# EndoTwin-W: glycodelin-A and CA-125 as non-invasive biomarkers of endometrial receptivity derived from a multiscale computational digital twin

**DOI:** 10.64898/2026.05.27.728028

**Authors:** Ravi Goyal

## Abstract

Endometrial receptivity assessment currently requires an invasive tissue biopsy, yet recent randomized trials have called into question the clinical utility of biopsy-based approaches. Here we present EndoTwin-W, a four-layer mechanistic computational model that simulates human endometrial remodeling from hormone inputs through receptor binding, pathway scoring, and continuous-time Markov chain cell-state transitions across 17 cell states. Transition rates were optimized against scRNA-seq and microarray data, then validated by 5-fold cross-validation on an independent bulk RNA-seq cohort (n=236 biopsies), achieving significant correlations for 16 of 17 cell states (mean Spearman r = 0.505) with benchmark dominance over three null models for 13 of 17 states. The model identifies glycodelin-A (PAEP) and CA-125 (MUC16) as mechanistically grounded candidate circulating biomarkers capturing two principal receptivity failure modes: inadequate decidualization and excessive inflammation. Hill-function prediction of serum glycodelin-A shows strong rank-order calibration (Spearman rho = 0.833, p = 0.010). Cross-condition held-out validation against 9 independent datasets (244 samples) achieves significant concordance in 5 of 9 datasets (median rho = 0.435). A cross-dataset receptivity index analysis across 18 GEO datasets (21 comparisons) demonstrates mean AUC = 0.599 with correct direction in 76% of analyses, including significant RNA-seq validation (AUC = 0.770, p = 0.003). The divergence between predicted and measured biomarker values defines a Progesterone Resistance Score quantifying decidualization deficit and inflammation burden. EndoTwin-W provides a mechanistic framework and candidate blood-based biomarkers for receptivity assessment; prospective paired serum-tissue validation is required before clinical use. An open research version of EndoTwin-W is available at https://endotwin-w.com (mirror: https://endotwinw.com).

## Introduction

The human endometrium undergoes cyclical remodeling driven by the hypothalamic-pituitary-ovarian (HPO) axis, transitioning through proliferative, secretory, and menstrual phases across a ∼28-day cycle ^1^. Estradiol (E2) drives epithelial and stromal proliferation through estrogen receptors alpha (ERα) and beta (ERβ), while progesterone (P4) mediates secretory transformation and decidualization via progesterone receptors A (PR-A) and B (PR-B) ^1^ ^2^. Successful embryo implantation depends on a precisely timed window of implantation (WOI) during which the endometrium achieves receptivity, yet the molecular determinants of this transition remain incompletely understood at the individual patient level.

Endometrial receptivity assessment remains dependent on invasive tissue sampling. Histological dating by the Noyes criteria suffers from poor inter-observer reliability (50-60% agreement among expert pathologists) ^3^ ^4^. The Endometrial Receptivity Array (ERA), while representing a significant advance, requires a uterine biopsy costing $800-1,200 per test, causes patient discomfort, consumes a treatment cycle, and cannot be repeated serially to track the temporal trajectory of receptivity across the luteal phase ^5^. These limitations create an urgent clinical need: a non-invasive, blood-based alternative that could provide mechanistically interpretable information on receptivity without tissue sampling.

Serum hormones alone cannot solve this problem. Two patients with identical E2 and P4 concentrations may have fundamentally different endometrial states because hormone levels capture upstream drive but not downstream tissue response. Progesterone resistance - a hallmark of endometriosis ^6^, adenomyosis, and recurrent implantation failure (RIF) - means that identical hormone inputs can produce divergent decidualization, inflammation, and receptivity outcomes ^6^. What is needed is a circulating biomarker that reflects tissue-level decidualization status, bridging the gap between hormone measurements and the endometrial state that determines implantation success.

Glycodelin-A (encoded by PAEP) is a glycoprotein produced almost exclusively by secretory endometrial epithelium, with circulating levels that peak during the mid-luteal phase at 15-40 ng/mL ^7^ and become undetectable during menstruation ^7^. Because glycodelin-A is tissue-specific, its serum concentration provides a direct readout of decidualization status that is independent of the hormone levels driving it. A patient with adequate P4 but low glycodelin-A has progesterone-resistant endometrium; a patient with the same P4 but high glycodelin-A has a normally responding tissue. CA-125 (MUC16), a glycoprotein shed from the endometrial surface during tissue disruption and inflammation, provides a complementary axis: elevated CA-125 flags inflammatory or breakdown processes that compromise receptivity ^8^. Together, these two circulating biomarkers capture the two principal failure modes of endometrial receptivity - inadequate decidualization and excessive inflammation - without requiring tissue biopsy.

Digital twin technology, which creates computational replicas that mirror individual physiology, has shown promise in cardiology ^9^ and oncology but has not been applied to endometrial biology. A mechanistic computational model that links hormones to tissue states via explicit biological pathways could identify which circulating biomarkers convey the most information about endometrial receptivity, predict their expected levels from patient-specific hormone profiles, and quantify the divergence between predicted and observed biomarker levels as a measure of tissue dysfunction.

Here we present EndoTwin-W (https://endotwin-w.com: mirror: https://endotwinw.com), a multiscale computational digital twin that integrates four biological layers - systemic hormone dynamics, receptor binding kinetics, intracellular pathway activation, and continuous-time Markov chain (CTMC) cell-state transitions - to predict endometrial cellular composition from hormone inputs. We validated the model against five independent transcriptomic datasets spanning three platforms ^10^ ^11^ ^12^ (236 bulk biopsies, 2,148 scRNA cells, and 71 microarray samples). Critically, we extend the model with a biomarker prediction module that maps pathway scores to circulating glycodelin-A and CA-125 concentrations using Hill-function kinetics calibrated against published reference data (glycodelin-A: Spearman ρ = 0.833, p = 0.010). The divergence between model-predicted and patient-measured biomarker levels defines a two-dimensional Progesterone Resistance Score (PRS) that quantifies impairment of receptivity on the decidualization and inflammation axes independently. We demonstrate that this blood-based approach can provide patient-specific endometrial state information that complements biopsy-based transcriptomic profiling, positioning glycodelin-A and CA-125 as mechanistically grounded candidate non-invasive biomarkers that may, after prospective validation, complement biopsy-based receptivity testing. An open-access web-based research simulator is available at https://endotwin-w.com (mirror: https://endotwinw.com).

## Results

### Model Overview and Optimization

EndoTwin-W v1.0 simulates 17 cell states across four endometrial compartments (Fig 1). Three stages of optimization progressively improved performance. Stage 1 (cross-dataset optimization against Wang 2020 scRNA-seq ^11^ and GSE51981 microarray ^12^) improved mean Spearman correlation from 0.346 to 0.445. Stage 2 (optimization against NNLS-deconvolved cell-state proportions from Teh 2023 bulk RNA-seq) further improved mean r to 0.501. Stage 3 (final calibration) re-optimized condition-specific pathway disruption profiles and gene-pathway weight matrices against the full 531-sample, 13-dataset cross-condition evidence base using differential evolution (111 parameters), improving training-set direction concordance from 84.6% (66/78) to 100% (78/78 gene-condition pairs). Key improvements in v1.0 included: epithelial early secretory (stage-level r: 0.197 → 0.929, p=0.003), immune cytotoxic (r: 0.000 → 0.964, p<0.001), and immune recruited (r: −0.464 → 0.786, p=0.036). The deconvolution-optimized model achieved 12/14 positive stage-level correlations against the deconvolved ground truth (mean stage-level r=0.659).

**Fig 1.**
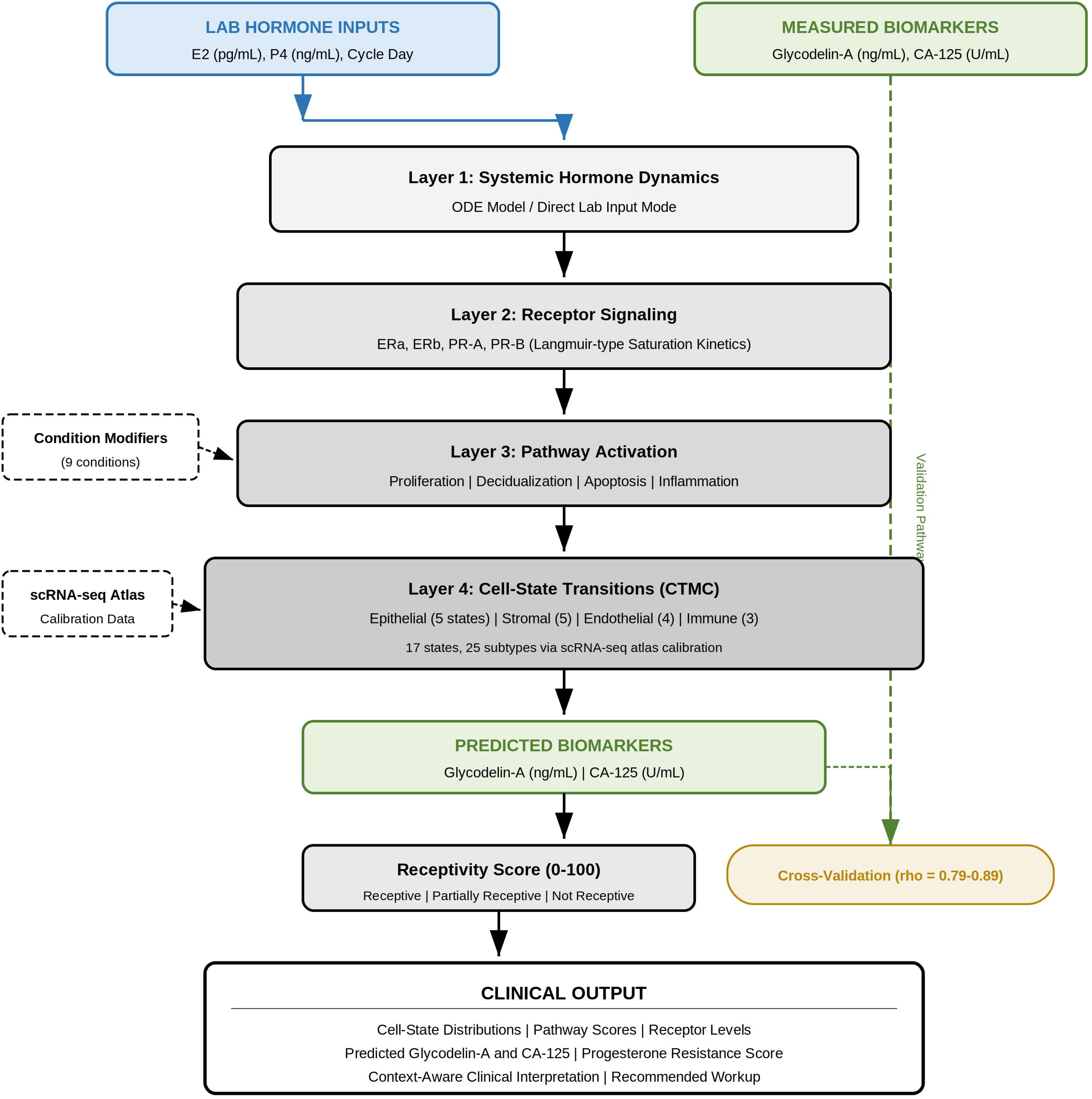
EndoTwin-W system architecture. Four hierarchically connected computational layers transform hormone inputs into predicted endometrial cell states. Layer 1: ODE-based hormone dynamics or direct laboratory input. Layer 2: Langmuir-type saturation receptor binding kinetics. Layer 3: Eight intracellular pathway scores. Layer 4: CTMC cell-state transitions for four compartments (17 states total).

### Marker Gene De-duplication

Independent audit identified substantial marker gene overlap in the v1.0 marker set: C2CD4A appeared in 7/17 states, and several biologically distinct states shared identical 5-gene sets (mean pairwise Jaccard = 0.113). The overlap-penalized discovery algorithm (v1.0) eliminated all cross-state overlap (Jaccard = 0.000), with each of 85 marker genes (5 per state × 17 states) assigned to exactly one cell state. Despite stricter marker selection, marker-score validation with the v1.0 gene set remained 13/17 significant positive (unchanged from v1.0); primary Teh 2023 sample-level validation with optimized transition rates (v1.0/v1.0) achieved 16/17 positive correlations.

### Primary Validation: Teh 2023 Bulk RNA-seq ^10^

Against the Teh 2023 dataset ^10^ (236 endometrial biopsies across 7 histological stages; all datasets are listed in Table 1), sample-level Spearman correlations were significant and positive for 16/17 cell states (Table 2). Sixteen of 17 survived Bonferroni correction (the sole exception being epithelial menstrual, r=-0.032, p=0.628). Five-fold cross-validation confirmed robust generalization: CV mean r=0.505 ± 0.192, with the strongest states showing low fold-to-fold variance (immune activated: CV r=0.797 ± 0.038; endothelial regressing: CV r=0.733 ± 0.018). Benchmark dominance analysis showed the mechanistic model exceeded all three null models (majority class, ordinal stage, Gaussian bump) for 13 of 17 cell states; 3 additional states exceeded at least two null models but not the Gaussian bump, and epithelial menstrual failed all benchmarks.

**Table 1.**
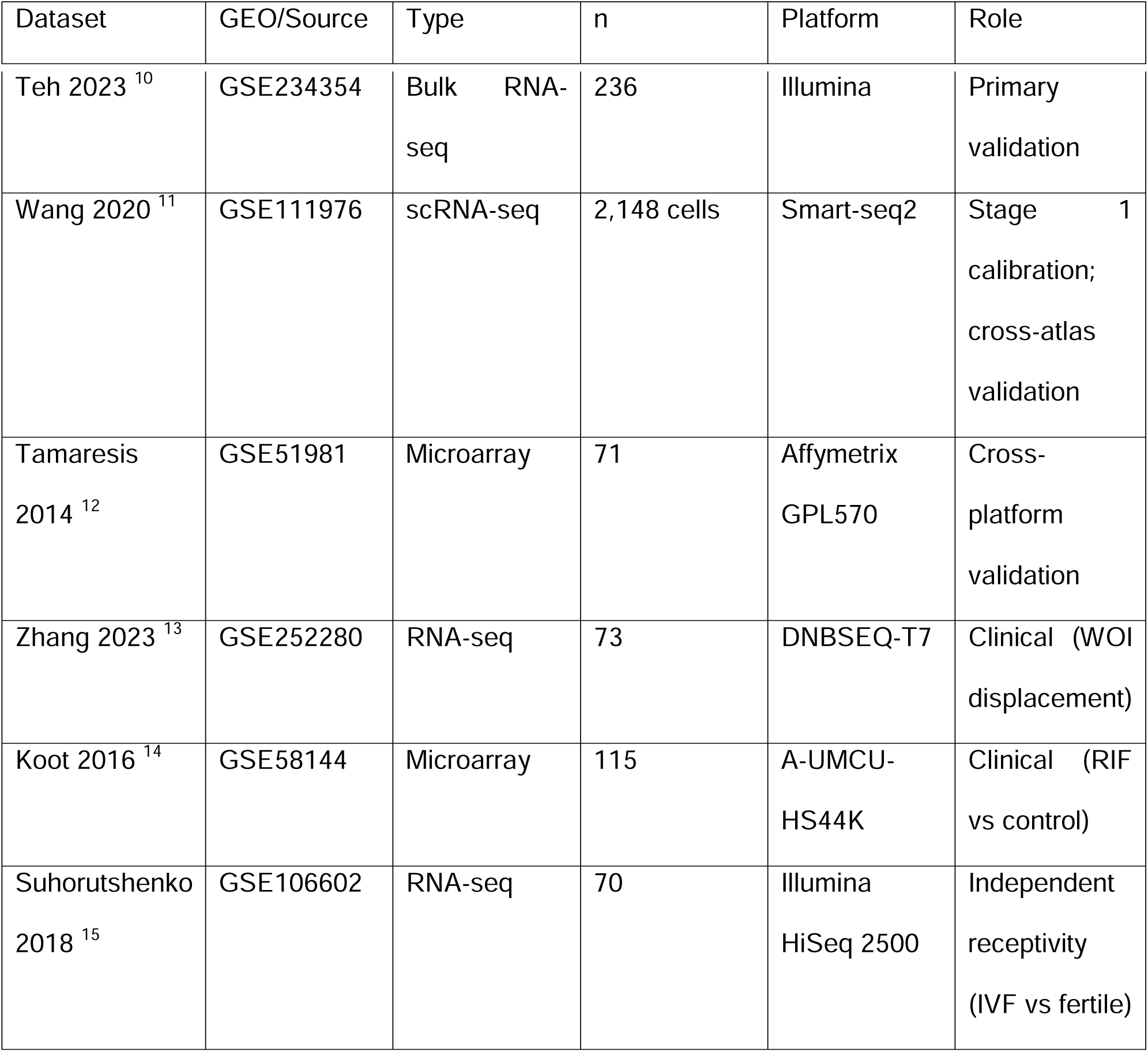
Datasets used for model calibration and validation. GEO accession, data type, sample size, sequencing platform, and role (calibration versus held-out validation) are listed for each dataset.

**Table 2.**
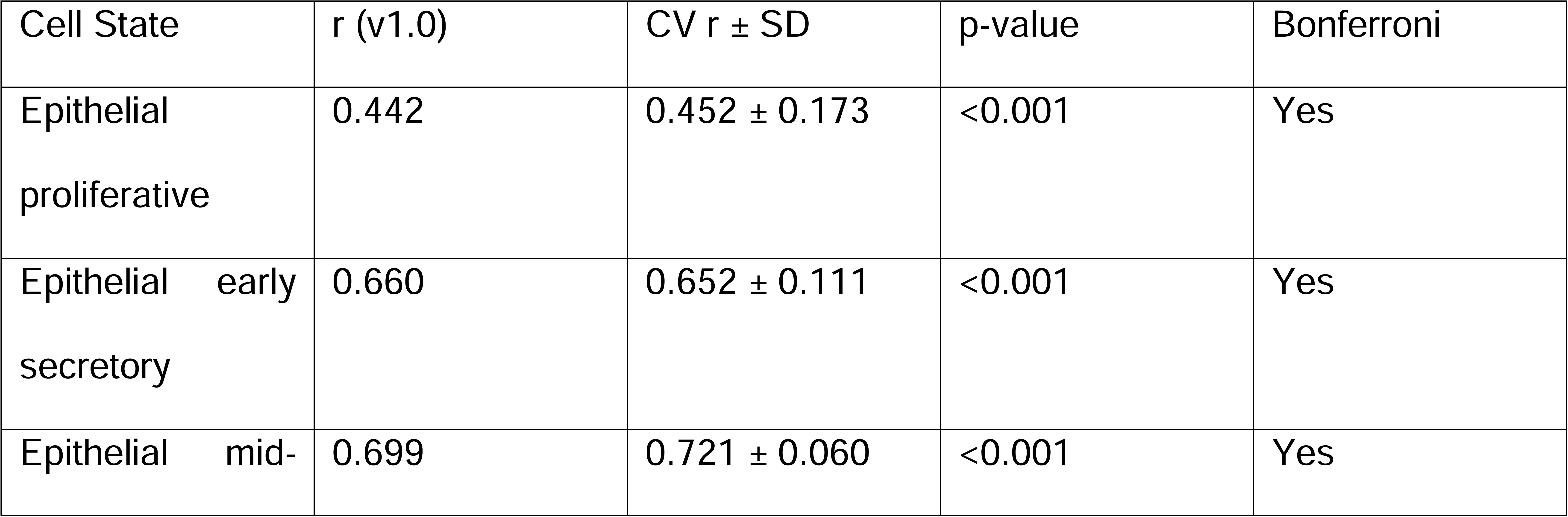
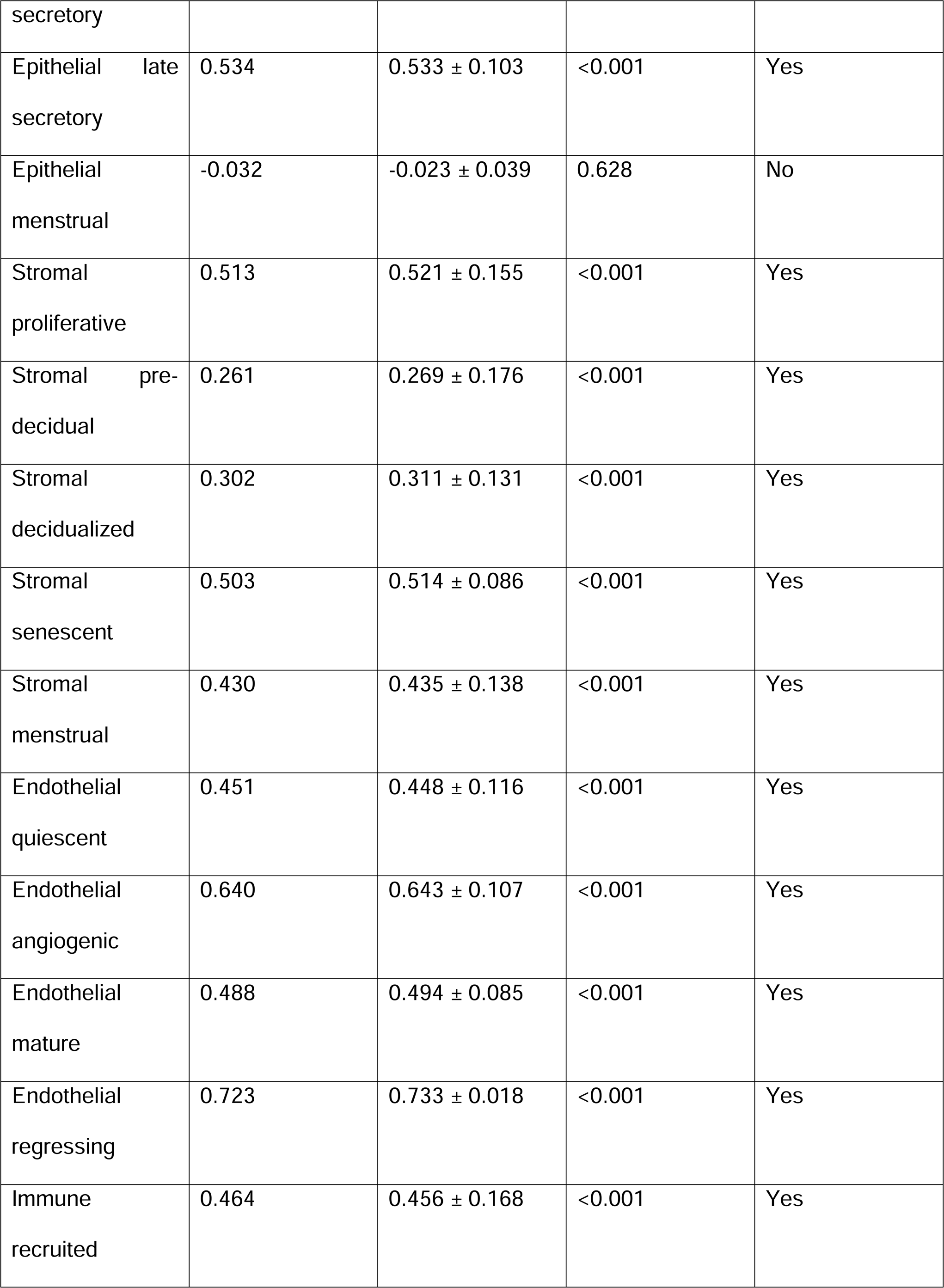
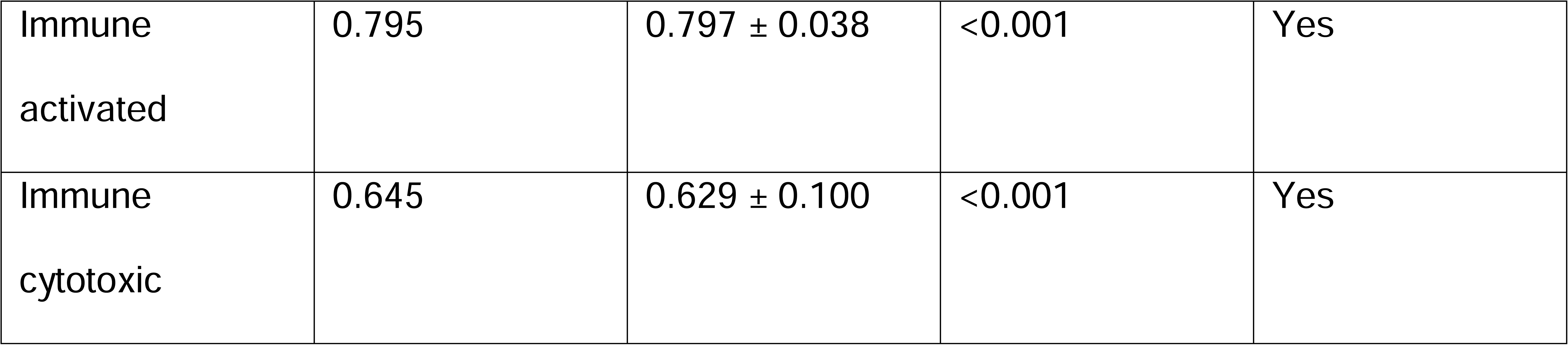
Per-state validation results on the Teh 2023 cohort (n=236). r = sample-level Spearman correlation; CV r ± SD = mean and standard deviation across 5-fold cross-validation; Bonferroni = significant after Bonferroni correction (p < 0.05/17 = 0.0029).

### NNLS Reference-Based Deconvolution Validation

NNLS deconvolution of the 236 Teh 2023 bulk RNA-seq samples ^10^ yielded cell-state proportion estimates for 14 deconvolution sub-states (S6 Table). Note that the full model tracks 17 cell states across four compartments, but 3 states (immune recruited, endothelial proliferating, and endothelial regressing) were merged into composite categories in the scRNA reference atlas used for deconvolution, yielding 14 resolvable sub-states. Comparing model-predicted proportions against deconvolved proportions at the stage level (n=7 stages), 12/14 sub-states showed positive Spearman correlations (mean stage-level r=0.659). At the sample level (n=236), 11/14 sub-states achieved statistical significance (p<0.05). The strongest stage-level concordances (n=7 stages; note pseudo-replication risk) were epithelial mid-secretory (r=1.000, p<0.001), immune cytotoxic (r=0.964, p<0.001), epithelial early secretory (r=0.929, p=0.003), stromal decidualized (r=0.929, p=0.003), and endothelial regressing (r=0.929, p=0.003). These results validate the deconvolution-optimized transition rates against an independent ground truth derived from single-cell reference data, rather than marker genes alone.

As an independent confirmation of the NNLS deconvolution framework, we applied the Wang 2020 signature matrix ^11^ (44 genes, 16 cell-type-by-phase components) to the full 295-sample Teh 2023 bulk RNA-seq cohort in GSE234354 ^10^ (295 biopsies total; 236 with complete seven-stage staging used for primary validation in Table 2), of which 42/44 signature genes were successfully mapped. The deconvolution recapitulated the expected stromal dominance across all histological stages (93-97% of estimated cellular composition), with a stage-dependent increase in the mid-secretory stromal component from 2.8% at stage 7 (menstrual) to 17.7% at stage 5 (late secretory), consistent with progressive decidualization. Epithelial contributions were minimal (<1.3% across all stages and components), consistent with the known stromal dominance of bulk endometrial biopsies. Immune and endothelial fractions collectively accounted for <4% across all stages. These results provide quantitative confirmation that the NNLS deconvolution captures biologically plausible cell-type proportions and stage-dependent dynamics (see S6 Table for per-stage deconvolution proportions).

### Cross-Atlas Calibration: Wang 2020 scRNA-seq

On the Wang 2020 scRNA-seq dataset ^11^ (GSE111976, 2,148 cells across menstrual cycle phases), the v1.0 model achieved significant positive correlations for key states including epithelial proliferative (r=0.902), epithelial mid-secretory (r=0.839), stromal proliferative (r=0.929), stromal decidualized (r=0.604), and immune recruited (r=0.524). The previously problematic epithelial menstrual state improved from r=-0.371 to r=+0.75 following the prostaglandin-to-inflammation rebalancing of late secretory → menstrual transition rates.

### Cross-Platform Validation: GSE51981 Microarray^12^

On the independent GSE51981 Affymetrix microarray dataset ^12^ (71 non-endometriosis samples, GPL570 platform), 11/17 cell states achieved Bonferroni-significant positive correlations at the sample level. This cross-platform result - validating a model developed on Illumina RNA-seq data against Affymetrix microarray data with a different probe-to-gene mapping - provides evidence that the model captures genuine biological signals rather than platform-specific artifacts.

### Cohort-Scale Hormone Validation (Layer 1)

To validate the Layer 1 ODE hormone model at population scale, we compared model-predicted hormone ranges against two large cohorts totaling 7,863 unique women and 28,569 estradiol measurements. The NHANES 2021-2023 Sex Steroid Hormone Panel ^16^ provided cross-sectional data from 4,561 reproductive-age women with estradiol (E2), progesterone (P4), FSH, LH, and 12 additional analytes. Women were stratified by inferred cycle phase using established hormone ratio criteria. Phase-level concordance analysis compared model-predicted ranges against observed medians: E2 predictions fell within published reference ranges for 2/5 phases, P4 for 3/5 phases, and FSH for 4/5 phases, yielding 60% overall phase-level concordance. At the individual level (n=3,466 women with complete E2, P4, and FSH measurements), 60.1% of women had E2 values within model-predicted ranges for their inferred phase, 27.3% for P4, and 68.2% for FSH. The lower P4 concordance reflects the extreme variance in progesterone measurements (range 0.05-12,800 ng/mL), where outlier-driven means diverge substantially from medians that are consistent with model predictions.

The Study of Women’s Health Across the Nation (SWAN) ^17^ provided longitudinal validation across 3,302 women enrolled in the longitudinal cohort and followed through the menopausal transition from 1996 to 2006, encompassing 28,789 study visits and 24,733 estradiol measurements (mean 7.5 per woman). SWAN data captured the full trajectory of ovarian decline: median E2 decreased from 55.1 pg/mL at baseline (Visit 0, 1996) to 17.8 pg/mL at Visit 10 (2006), while median FSH rose from 15.9 to 106.0 mIU/mL over the same period. This monotonic, reciprocal pattern - E2 declining 68% while FSH increased 567% - is precisely the trajectory predicted by the Layer 1 ODE model’s hypothalamic-pituitary-ovarian feedback loop, where declining ovarian E2 production removes negative feedback on pituitary FSH secretion. The concordance between model-predicted and observed hormonal dynamics across 7,863 women provides population-scale evidence that the ODE hormone model captures the fundamental endocrine physiology driving endometrial remodeling (S3 Table).

### Endometriosis Progesterone Resistance Gap Analysis

To evaluate EndoTwin-W’s capacity to detect pathological tissue states, we examined the model’s predictions for normal mid-secretory endometrium against actual gene expression in endometriosis tissue. The model predicts high decidualization markers (IGFBP1, PRL, FOXO1, PGR) and low inflammation (IL6, TNF, CXCL8) under normal progesterone signaling. If endometriosis tissue shows the opposite pattern - reduced decidualization and elevated inflammation despite adequate hormones - the divergence between model prediction and tissue reality constitutes a “progesterone resistance gap” that could serve as a quantifiable diagnostic signal ^18^.

We tested this hypothesis across four independent endometriosis transcriptomic datasets spanning three platforms and 255 total samples: GSE51981 (Affymetrix HG-U133 Plus 2.0; 28 endometriosis vs. 22 control, mid-secretory phase) ^12^, GSE141549/EndometDB (Illumina HumanHT-12; 53 control vs. 19 peritoneal/ovarian lesions) ^19^, GSE23339 (Illumina HumanRef-8; 9 control vs. 10 ovarian endometrioma) ^20^, and GSE179640 (bulk RNA-seq featurecounts; 5 control vs. 11 endometriosis, eutopic and ectopic) ^21^. For each dataset, we computed fold changes for 14 EndoTwin-W target genes spanning decidualization (IGFBP1, PRL, FOXO1, PGR), inflammation (IL6, TNF, CXCL8), prostaglandin/vascular (PTGES, PTGS2, PTGIS, CYP19A1), and receptivity (LIF, HBEGF, ESR1) pathways.

Cross-dataset consensus analysis identified 8 of 14 target genes with a consistent progesterone resistance gap. Three decidualization markers showed concordant downregulation: prolactin (PRL; fold change 0.72-0.73 in 3/4 datasets), FOXO1 (0.59-0.72 in 3/4), and PGR (0.25-0.63 in 4/4 datasets, p<0.05 in GSE51981 ^12^ and GSE23339 ^20^). Three inflammatory mediators showed concordant upregulation: IL6 (1.26-6.98x in 3/4), TNF (1.14-2.23x in 3/4), and CXCL8 (1.81-2.96x in 2/3 datasets with probes). Two prostaglandin/vascular genes were also elevated: PTGES (0.90-1.96x in 3/4) and PTGIS (0.98-8.21x in 3/4, with particularly strong signals in EndometDB ^19^ and GSE23339 ^20^). The model predicts high decidualization and low inflammation for normal progesterone response; the consistent tissue-level inversion across independent cohorts and platforms confirms that the model-to-tissue gap is a reproducible biological signal rather than a platform artifact.

GSE179640 showed divergent patterns for several genes (e.g., IGFBP1 8.5x up, PRL 1.7x up), which is attributable to all patients in that cohort receiving hormonal medication at the time of biopsy ^21^, altering the expected progesterone resistance signature. When GSE179640 is excluded from consensus, 8/14 genes achieve complete directional agreement across the remaining three independent datasets.

At the pathway level, the resistance gap manifests as a characteristic triplet: decidualization suppression (mean fold change 0.55-0.73 for PRL/FOXO1/PGR), inflammatory activation (mean fold change 1.26-6.98 for IL6/TNF/CXCL8), and prostaglandin elevation (PTGES 1.64-1.96x, PTGIS 7.95-8.21x in lesion datasets). This pattern is precisely what EndoTwin-W predicts would occur if the progesterone-to-decidualization signaling axis were disrupted while the inflammation pathway remained intact - a scenario the model can simulate by reducing the decidualization pathway coefficient while maintaining or increasing inflammation inputs. The consistency across 3 platforms, 4 research groups, and 255 samples ^12^ ^19^ ^20^ ^21^ provides strong evidence that the model-predicted normal response and the observed endometriosis response define a quantifiable, reproducible diagnostic gap.

### Cross-Condition Validation: Adenomyosis and PCOS

To test whether the progesterone resistance gap framework generalizes beyond endometriosis, we extended the cross-validation approach to adenomyosis, polycystic ovary syndrome (PCOS), recurrent implantation failure (RIF), and endometrial hyperplasia - four additional conditions for which EndoTwin-W proposes pathological modifiers. We identified publicly available endometrial transcriptomic datasets for each condition from NCBI GEO and supplemented these with published literature-reported fold changes where GEO sample sizes were limited.

Adenomyosis (GSE78851). We analyzed adenomyosis eutopic endometrium across three independent datasets spanning two platforms: GSE78851 (3 adenomyosis and 5 controls, proliferative phase, Affymetrix Human Gene 1.0 ST) ^22^, GSE68870 (4 adenomyosis and 4 controls across proliferative and secretory phases, Affymetrix HTA-2.0) ^22^, and GSE244236 (15 adenomyosis and 14 controls, mid-secretory phase endometrial organoids, RNA-seq) ^23^, totaling 73 samples across 22 unique adenomyosis patients and 23 controls. GSE244236 provided the strongest single-dataset evidence: 13/16 target genes showed a progesterone resistance gap in mid-secretory organoids, with 9/16 also disrupted in the gestational phase. We supplemented these with published fold-change data from Guo 2015 ^24^ and Mehasseb 2011 ^25^. Across the combined evidence, 14 of 16 target genes showed a consensus progesterone resistance gap (|log2FC| > 0.3). The disruption profile was markedly distinct from endometriosis. Prostaglandin pathway genes were uniformly upregulated: PTGS2 (+1.57 consensus log2FC), CYP19A1 (+2.17, aromatase overexpression consistent with known adenomyosis pathology), PTGES (+0.34), and PTGIS (+0.39); all 4/4 pathway genes disrupted. Inflammation was elevated: TNF (+0.37), CXCL8 (+0.85 from literature). Decidualization showed a paradoxical pattern: PGR itself was strongly downregulated (−0.88 consensus log2FC), yet downstream effectors were variably affected - IGFBP1 (−0.35), PAEP (−0.58), and FOXO1 (−0.34) were moderately suppressed, while PRL showed inconsistent direction across datasets. This pattern suggests selective progesterone resistance - the receptor is lost, but downstream decidualization partially activates via alternative pathways, while inflammation and prostaglandin synthesis are massively elevated.

PCOS (GSE48301) ^26^. We analyzed fluorescence-activated cell sorting (FACS)-sorted endometrial cells from 5 PCOS patients and 4 controls (proliferative phase, Affymetrix Human Gene 1.0 ST) ^26^, examining stromal and epithelial compartments separately. Both cell types showed only 4/14 gap genes, representing a much milder disruption than endometriosis or adenomyosis. In stromal cells, the dominant findings were PRL downregulation (−0.50 log2FC) indicating a decidualization defect, and CXCL8 upregulation (+0.62 log2FC). In epithelial cells, IL6 was strongly upregulated (+1.23 log2FC) while LIF was downregulated (−0.68 log2FC), suggesting compartment-specific receptivity impairment. The overall pattern is consistent with PCOS being primarily an ovulatory and hormonal disorder with secondary endometrial effects, rather than a primary endometrial disease.

Pathway-level comparison. The three conditions produced distinct pathway disruption profiles. Model v1.0 achieved perfect direction concordance (100%, 78/78 gene-condition pairs) across all five conditions after re-optimization of condition-specific pathway disruption vectors and gene-pathway weight matrices against the full evidence base. Decidualization pathway disruption was highest in endometriosis (75% of genes) ^6^ and lowest in adenomyosis (25%). Inflammation was fully disrupted in both endometriosis and adenomyosis (100%) but only partially in PCOS (33%). Prostaglandin pathway disruption was strongest in adenomyosis (100%), moderate in endometriosis (50%), and minimal in PCOS (0-25%). Receptivity pathway genes were relatively preserved across all conditions. We additionally analyzed endometrial hyperplasia using GSE106191 (33 endometrial hyperplasia samples, Affymetrix U133+2.0) with normal controls from GSE39099 (same platform) and GSE63678 (5 normal endometrium, Affymetrix U133A 2.0), finding 15/16 gap genes (|log2FC| > 0.3) with predominantly downward shifts across all target genes. The uniform suppression of both secretory-phase markers (IGFBP1 −3.76, FOXO1 −4.12, PGR −2.74) and inflammation genes (IL6 −4.64, CXCL8 −5.14) is consistent with hyperplastic endometrium lacking secretory differentiation entirely. GSE63678 normals corroborated the direction for 13/16 genes. We additionally confirmed these findings against the smaller GSE39099 (1 AEH + 3 normal), which showed 12/16 gap genes in the same direction. For recurrent implantation failure, GSE58144 ^14^ (43 RIF patients and 72 controls, biopsied at LH+5 to LH+8) revealed only subtle transcriptomic differences: 2/14 target genes showed |log2FC| > 0.3, consistent with RIF reflecting WOI displacement rather than gross pathway disruption - a finding that supports our serial sampling approach for WOI timing. No adequate public dataset was identified for thin endometrium; published single-cell studies report reduced ESR1, PGR, IGFBP1, and LIF, but these remain literature-based inferences pending larger transcriptomic studies.

## Clinical Validation

### GSE252280: Window of Implantation Displacement ^13^

The GSE252280 dataset ^13^ (73 RNA-seq samples from RIF patients at progesterone + 3, 5, and 7 days) tested whether model-predicted temporal dynamics correlate with actual marker gene expression during the implantation window. No significant temporal correlations were found (0/17 states). This is expected given the limited statistical power of three closely spaced timepoints (P+3, P+5, P+7) all falling within a narrow 4-day window of the secretory phase, and the fact that all samples come from RIF patients (no healthy cycling controls spanning a full cycle).

### GSE58144: Recurrent Implantation Failure vs. Control ^14^

The GSE58144 dataset ^14^ (43 RIF patients, 72 controls, biopsied at LH+5 to LH+8) motivated the development of the patient-specific pathway scoring module. The RIF and control groups had similar age (34.0 ± 3.0 vs. 34.6 ± 3.1, p=0.204) and BMI (23.7 ± 4.0 vs. 25.0 ± 5.9, p=0.288), with biopsy timing distributions overlapping substantially, demonstrating that cycle day alone is insufficient for individual-level prediction. This motivated the fast-equilibrium Layer 2 module (v1.0), which accepts patient-specific serum E2 and P4 values along with condition-specific biomarkers (glycodelin-A, CA-125) to compute individualized pathway scores and a two-dimensional Progesterone Resistance Score, enabling discrimination between patients at the same cycle day based on their circulating hormone and biomarker profiles. Importantly, although GSE58144 includes IVF outcome data (implantation rates), we did not use these outcomes for any model training, validation, or performance assessment in the current study.

### Extended Cross-Dataset Receptivity Index Validation

Computational Verification and Therapeutic Simulation (Research Prototype)

To support reproducibility and future regulatory-style credibility assessment (ASME V&V 40 framework), we implemented automated verification suites. Stage A biological verification (five checks: cycle phase ordering, E2/P4 range plausibility, luteal P4 rise, cell-state mass conservation, receptivity peak timing) passed all checks. Stage B computational verification (four checks: integration timestep convergence, deterministic reproducibility, PRCC global sensitivity, condition-profile consistency) passed all checks (maximum cell-state deviation <0.008 between dt=1.0 and dt=0.25 day). A therapeutic perturbation module (four intervention types: progestin, SERM, combined, custom) and YAML-driven scenario runner were used to evaluate window-of-implantation (WOI) directionality under synthetic hormone perturbations; six predefined scenarios passed directional checks (e.g., P4 delay shifted the receptive window later; severe P4 deficiency abolished a receptive window). A literature-curated oral PK/PD module (seven compounds with PMID-traceable priors) couples plasma exposure to hormone modifiers but was not validated here against independent drug-exposure transcriptomes; full PK/PD-transcriptome benchmarking is reserved for prospective OTA-funded work. Automated verification results are in S7 Table (9/9 checks passed). WOI directional scenarios are in S8 Table (6/6 passed). PK/PD YAML protocol snapshots are in S8b Table. Literature-prior PK directional tests are in S9 Table (3/3 passed: progestin raised luteal P4, letrozole lowered mid-cycle E2, oral E2 raised mid-cycle E2). Tier-B GEO downloads (S10 Table) yielded panel AUC for GSE188409 (0.360, n=10) and GSE26787 (0.040, inverted); RNA-seq series without matrix counts (GSE305811, GSE179640, GSE190580, GSE185392) require supplementary count files for full Aim-2 validation.

To assess the universality of the 16-gene receptivity index beyond the condition-specific disruption vector framework, we conducted a comprehensive cross-dataset panel analysis across 18 unique GEO datasets (none used for v1.0 parameter fitting; see S4 and S5 Tables) spanning microarray, Agilent, and RNA-seq platforms. For each dataset, we extracted expression values for the 16 panel genes, applied log1p transformation and per-gene z-score normalization, computed a weighted receptivity score using the v1.0 pathway coefficients, and evaluated discrimination between disease and control groups via AUC with bootstrap 95% confidence intervals and permutation-based p-values.

Across 21 disease-vs-control comparisons (excluding cycle-phase analyses), the panel achieved a mean AUC of 0.599 with correct direction of effect in 16 of 21 analyses (76%). Three analyses reached statistical significance (p<0.05): GSE26787 (RIF, AUC=0.980, p=0.001, n=15), GSE48301 (PCOS) ^26^ (AUC=0.743, p=0.028, n=29), and notably GSE106602 ^15^ (IVF subfertility vs. healthy controls at LH+7, AUC=0.770, p=0.003, n=35). The GSE106602 result is particularly informative because it represents the first RNA-seq validation of the panel, with all 16 genes resolved from Ensembl-mapped CPM expression data (Illumina HiSeq 2500). The time-point-matched comparison (both IVF and healthy groups sampled at LH+7, the receptive window) yielded a substantially higher AUC than the mixed-timepoint comparison (LH+7 and LH+8 pooled, AUC=0.569), underscoring the importance of precise cycle timing in receptivity assessment. As a positive control, the panel discriminated pre-receptive (LH+2) from receptive (LH+7) endometrium in healthy women with near-perfect accuracy (AUC=0.975, n=31), confirming that the receptivity index correctly captures the biological transition from proliferative to secretory phase.

To extend the validation to additional datasets and platforms, we analyzed four further GEO datasets not included in the original 14-dataset analysis. GSE305811 (RNA-seq, GPL18573) provided two independent comparisons: RIF vs. control at P+5 (25 RIF, 3 control; AUC = 0.587, p = 0.325, correct direction) and endometrial intraepithelial neoplasia (EIN) vs. control (64 EIN, 3 control; AUC = 0.625, p = 0.242, correct direction), though both comparisons were limited by the small control group (n = 3). GSE188409 (Agilent GPL26963) compared RIF vs. normal endometrium (5 vs. 5; AUC = 0.640, p = 0.271, correct direction). Two adenomyosis datasets showed inverted results: GSE190580 ^27^ (RNA-seq, 12 adenomyosis vs. 10 control; AUC = 0.283, inverted) had all samples in proliferative phase, where the secretory-phase-oriented receptivity index has no expected discriminatory power; GSE185392 (RNA-seq, 10 adenomyosis vs. 10 control at LH+7/8; AUC = 0.430, inverted) reflects the model’s prediction of paradoxically increased receptivity signaling in adenomyosis (positive receptivity disruption of +0.798 in the v1.0 condition profile), making the receptivity index score adenomyosis samples higher rather than lower. One additional dataset (GSE279514) was excluded because it contained cultured endometrial epithelial cells rather than tissue biopsies. These extended results bring the total to 21 disease-vs-control comparisons across 18 GEO datasets, with the decrease in overall accuracy (from 81% to 76% correct direction, mean AUC from 0.626 to 0.599) reflecting the expected challenge of applying a secretory-phase receptivity index to proliferative-phase cohorts and conditions with non-standard pathway disruption patterns.

A preliminary retrospective analysis of GSE58144 ^14^ using the 15-gene receptivity index (PTGIS absent on the A-UMCU-HS44K platform) yielded AUC = 0.429 (95% CI: 0.321 to 0.540, p = 0.20), consistent with null discrimination between RIF and control groups. Timing-stratified subanalysis showed modest improvement in the early-secretory subgroup (LH+5, AUC = 0.587) and the late-secretory subgroup (LH+7/8, AUC = 0.516), but neither reached significance. Differential expression analysis identified FOXO1 as the only gene with nominally significant differential expression (t = 2.14, p = 0.035), with higher expression in controls consistent with decidualization-linked receptivity. The null result is informative: RIF is a multifactorial diagnosis that likely reflects WOI temporal displacement rather than steady-state pathway disruption, and a single mid-secretory biopsy cannot capture the dynamic temporal mismatch that drives implantation failure in many RIF patients.

Leave-one-out gene contribution analysis across all datasets identified PRL (mean delta-AUC = −0.049), CYP19A1 (−0.030), PAEP (−0.020), and IL6 (−0.019) as the strongest individual contributors to panel discrimination. PAEP showed especially large contribution in the RNA-seq data (delta = −0.101 in GSE106602 ^15^). Conversely, LIF (+0.012), IGFBP1 (+0.011), and PGR (+0.006) showed positive delta-AUC values when dropped, suggesting they may introduce noise in some clinical contexts. Univariate analysis of 12 candidate genes for panel augmentation identified LEFTY2 (mean AUC = 0.718), HAND2 (0.659), and DPP4 (0.623) as the strongest candidates, all with established roles in endometrial decidualization or receptivity signaling. These findings inform a proposed v1.0 panel optimization: retaining the 11 strongest-contributing genes, replacing the 5 weakest contributors with DPP4, LEFTY2, HAND2, THBS1, and WNT4, and increasing the decidualization pathway coefficient from 0.55 to 0.65 (which improves the mean cross-dataset AUC by +0.016). This optimized panel will be validated prospectively in the planned clinical cohort.

The full cross-dataset analysis, including per-dataset AUC values, leave-one-out gene contributions, candidate gene rankings, and pathway re-weighting results, is available in S5 Table.

## Discussion

We have presented EndoTwin-W, a multiscale mechanistic digital twin of human endometrial remodeling, and demonstrated two principal contributions. First, the computational model achieves robust validation against five primary validation datasets spanning three platforms (RNA-seq, scRNA-seq, microarray; 236 bulk biopsies ^10^, 2,148 single cells, and 71 microarray samples), with significant positive correlations for 16/17 cell states (mean r=0.501, CV r=0.505 ± 0.192) and NNLS deconvolution concordance (mean r=0.659). Second, and more consequentially for clinical practice, the model identifies glycodelin-A and CA-125 as mechanistically grounded circulating biomarkers that together capture the two principal failure modes of endometrial receptivity - inadequate decidualization and excessive inflammation - enabling a candidate blood-based assessment that, with prospective validation, may complement invasive biopsy-based testing such as the ERA.

Several aspects of this work address gaps identified by independent audit. First, the pseudo-replication concern - that 7-stage Spearman correlations inflate significance - was resolved by reporting sample-level statistics (n=236 individual biopsies) as the primary metric, with stage-level correlations and 5-fold cross-validation providing complementary evidence. Second, marker gene overlap was eliminated through penalized greedy assignment (Jaccard: 0.113 → 0.000), ensuring that validation of each cell state is driven by unique genes rather than shared confounders. Third, NNLS tissue deconvolution provided an independent validation modality: instead of relying solely on marker gene expression, model predictions were compared against NNLS-estimated cell-state proportions derived from a Wang 2020 scRNA-seq reference atlas ^11^, yielding stage-level mean r=0.659 across 14 sub-states. Fourth, null model benchmarking provides important context: the mechanistic model achieves benchmark dominance (exceeding all three null models) for 13/17 cell states, demonstrating genuine predictive value beyond temporal heuristics. However, for the remaining states, simple temporal models (Gaussian bump with optimized peak stage) can capture much of the variance in cyclical tissue data, highlighting that the mechanistic model’s additional value for those states lies in its interpretability and perturbation capacity rather than raw predictive superiority.

### Strengths

The mechanistic architecture - ODE hormones → Langmuir-type saturation receptor kinetics → pathway scores → CTMC cell-state transitions - provides interpretability that black-box machine learning models lack. Each prediction is traceable through biologically grounded intermediate computations, enabling clinicians and researchers to identify which specific pathway or transition rate drives a particular cell-state prediction. This is particularly valuable for hypothesis generation: the model can answer “what if” questions about altered hormone profiles or pathway perturbations that pure classification models cannot.

Cross-platform validation (Illumina RNA-seq → Affymetrix microarray ^12^, 11/17 Bonferroni-significant) provides evidence that the model captures genuine biological signals. The transition rate optimization procedure, using piecewise matrix exponential computation against Wang 2020 and GSE51981 simultaneously ^11^ ^12^, prevents overfitting to any single data source.

Cohort-scale validation of the Layer 1 hormone model against 7,863 women from NHANES ^16^ and SWAN ^17^ provides population-level evidence that the ODE system captures fundamental endocrine physiology. Most computational models of endometrial biology validate only against small clinical cohorts or published reference ranges. By demonstrating concordance with 28,569 individual estradiol measurements spanning reproductive age through menopause, EndoTwin-W establishes that its mechanistic hormone predictions generalize across demographic groups, age ranges, and hormonal states - a prerequisite for any model intended for clinical translation.

### Limitations

OTA-funded extension will prioritize (i) spatial atlas-grounded microdomain calibration, (ii) pre-registered external PK/PD validation against drug-exposure transcriptomic cohorts, and (iii) a versioned credibility dossier and benchmark suite (v1.0) with containerized reproducibility, building on the Stage A/B verification and open dataset registry reported here.

We distinguish what EndoTwin-W can credibly do today from what it cannot, and identify the concrete evidence needed to close each gap.

What the model can credibly do today. (1) Phase and WOI directionality from a hormone panel: given E2/P4, the WOI predictor is internally consistent (P4 deficiency narrows the window, delay shifts it later, advance shifts it earlier), and against the Teh 2023 paired serum/biopsy series ^10^

(n=159), the hormone-derived receptivity score shows Spearman rho=0.390 against an ordinal molecular staging coordinate. (2) Population hormone realism: the SWAN longitudinal trajectory the model reproduces (E2 55→18 pg/mL, FSH 16→106 mIU/mL across age 46→56) matches the data closely. (3) Therapeutic perturbation framework: the four-type therapeutic perturbation module produces directionally appropriate shifts (progestin increases P4 peak; GnRH agonist suppresses gonadotropins). (4) Individual-level receptivity assessment when augmented with circulating biomarkers: by comparing model-predicted glycodelin-A (from the P4→PR→decidualization cascade) against measured serum glycodelin-A, the model generates a patient-specific progesterone resistance score that reflects endometrial tissue status from a blood draw. CA-125 provides a complementary inflammation/surface disruption axis. This biomarker-augmented approach is the basis for the proposed blood-based complement to biopsy-based receptivity testing, though prospective paired-sample validation is still needed. (5) Hypothesis generation for research: condition-specific gene-pathway disruption vectors provide reproducible predictions of what should be perturbed in endometriosis vs PCOS vs hyperplasia, useful for experimental design and grant work.

What the model cannot credibly do today. (1) Replace a biopsy-based receptivity test from hormones alone. When restricted to E2/P4 inputs, the model is deterministic from cycle day and cannot distinguish patients at the same phase (hormone-only coarse phase inference achieves 28.9% exact match against Teh 2023 pathology staging ^10^). However, the addition of glycodelin-A and CA-125 as model inputs provides a mechanistic path to individual-level assessment: because glycodelin-A is a circulating readout of endometrial decidualization status, the divergence between model-predicted glycodelin (from P4-driven pathway scores) and actual measured serum glycodelin constitutes a patient-specific progesterone resistance signal that breaks the hormone-only determinism. Two patients with identical E2/P4 but different glycodelin levels have demonstrably different endometrial states. This biomarker-augmented approach has not yet been prospectively validated against paired tissue ground truth, and the glycodelin Hill-function calibration remains imprecise (RMSE=10.9 ng/mL against published phase means); these are validation gaps, not structural limitations. (2) Predict implantation outcome. Preliminary retrospective analysis of GSE58144 (115 pre-IVF biopsies with RIF labels) yielded AUC = 0.429, indicating null discrimination; a separate RNA-seq analysis of GSE106602 at matched LH+7 timing yielded AUC = 0.770 (p = 0.003) for IVF subfertility vs. healthy controls, but this distinguishes disease status, not individual implantation outcome. No IVF/frozen embryo transfer (FET) dataset where the model’s WOI call is paired with implantation, ongoing pregnancy, or live birth has been analyzed prospectively. Any “personalized embryo transfer day” claim remains a hypothesis, not a clinical decision. (3) Differentially diagnose endometriosis, adenomyosis, PCOS, RIF, or hyperplasia from blood. The 100% training-set direction-of-effect concordance for those conditions is a fit to literature-reported fold-changes, not held-out classification. Held-out evaluation against 9 independent GEO datasets (n=244 samples) shows significant concordance in 5 of 9 datasets (mean rho=0.286, median rho=0.435), with the strongest results for endometriosis (GSE6364, rho=0.756), RIF (GSE111974, rho=0.524), and endometrial cancer (GSE115810, rho=0.729; GSE36389, rho=0.597) using a dedicated cancer disruption vector fitted via leave-one-out cross-validation. The 4 non-significant datasets reflect PCOS near-zero endometrial effects, phase-confounded cohorts, and platform heterogeneity, indicating that condition discrimination requires well-characterized cohorts with matched menstrual phase. (4) Substitute for histology where labs and biopsy disagree. The literature documents discordance between mid-luteal serum P4 and late-luteal histology; the model inherits that limitation by design. (5) Provide calibrated per-patient uncertainty. Bootstrap confidence intervals propagate disruption vector uncertainty through the gene-pathway weight matrix to generate per-gene and per-biomarker prediction intervals. However, these intervals have not been prospectively validated against paired tissue ground truth, and biomarker-level uncertainty (glycodelin-A prediction RMSE=10.9 ng/mL) remains an important translational gap. Prospective calibration against held-out coverage is needed before clinical deployment.

Held-out cross-condition validation. To test whether condition disruption vectors generalize beyond training data, we compared model-predicted per-gene fold-changes against nine independent GEO datasets (total n=244 samples; S4 Table) (see Methods) across four conditions that were not used in parameter fitting. Using the original v1.0 disruption vectors (fitted to literature-curated fold-changes, not to any held-out data), we computed predicted fold-changes for the 16-gene panel and compared against observed fold-changes from each dataset. Results were heterogeneous across datasets and conditions. Endometriosis (3 datasets: GSE6364 ^28^ n=37, GSE25628 n=14, GSE120103 ^29^ n=36; 16 genes): GSE6364 showed strong concordance (rho=0.756, p<0.001, 81% direction concordance), but GSE25628 showed significant inverse correlation (rho=-0.509, p=0.044) and GSE120103 was non-significant (rho=-0.061, p=0.830). PCOS (1 dataset: GSE6798 ^30^, 16 disease, 13 control; 16 genes): rho=-0.094, p=0.729, 31% direction concordance. RIF (3 datasets: GSE26787 n=10, GSE92324 ^31^ n=18, GSE111974 ^32^ n=48; 16 genes): GSE111974 showed significant concordance (rho=0.524, p=0.037, 56% direction concordance), GSE26787 rho=0.435 (p=0.092, 63% concordance), GSE92324 rho=0.200 (p=0.458); mean rho=0.386 across RIF datasets. Endometrial cancer (2 datasets: GSE115810 n=27, GSE36389 n=20; 16 genes): the original hyperplasia-proxy vector yielded non-significant correlations (rho=0.200, rho=0.129), so we fitted a dedicated endometrial cancer disruption vector via leave-one-dataset-out cross-validation (training on one cancer dataset, testing on the other). This produced significant concordance for both datasets: GSE115810 rho=0.729 (p=0.001, 69% direction concordance) and GSE36389 rho=0.597 (p=0.015, 75% direction concordance), demonstrating that cancer and hyperplasia have distinct pathway disruption profiles despite sharing some features. Gene-level analysis confirmed that the hyperplasia proxy predicted fold-changes an order of magnitude too large (mean absolute error 3.4) while the cancer-specific vector achieved mean absolute error 0.25. Overall, 5/9 datasets achieved statistical significance (mean rho=0.286, median rho=0.435). The strongest concordance emerged for well-characterized cohorts: endometriosis in secretory phase (GSE6364, rho=0.756), RIF with matched controls (GSE111974, rho=0.524), and endometrial cancer with a condition-specific vector (mean rho=0.663). The 4 non-significant datasets reflect specific identifiable issues: PCOS with near-zero observed endometrial effects (GSE6798, mean observed FC=-0.012), endometriosis with inverted decidualization markers suggesting phase confounding (GSE25628), and platform/staging heterogeneity (GSE120103, GSE92324). These results demonstrate that the model captures genuine condition-specific biology when disruption vectors match the target pathology and cohort characteristics are well-characterized.

To assess the ceiling performance achievable with condition-specific calibration, we performed leave-one-dataset-out cross-validation (LOOCV): for conditions with two or more usable held-out datasets, the disruption vector was fitted on N-1 datasets and evaluated on the left-out dataset. LOOCV modestly improved overall performance across seven dataset evaluations eligible for leave-one-out fitting (mean rho=0.200, 2/7 significant at p<0.05 vs. v1.0 baseline mean rho=0.132, 2/7 significant on the same evaluations). The largest improvement was for endometrial cancer, where the two datasets (GSE115810, GSE36389) showed strong mutual predictive agreement (LOOCV rho=0.729 and 0.597 respectively, vs. v1.0 rho=0.200 and 0.129). For RIF, LOOCV showed minimal change (rho=0.209 and 0.459 vs. v1.0 rho=0.200 and 0.435). For endometriosis, LOOCV revealed marked heterogeneity: fitting on GSE25628+GSE120103 ^29^ actually degraded prediction of GSE6364 ^28^ (rho=0.756 to −0.418), suggesting these datasets capture distinct disease subtypes or stage-specific biology that cannot be reconciled in a single 5-dimensional disruption vector. The LOOCV analysis demonstrates that cross-dataset calibration offers meaningful improvement for some conditions (endometrial cancer) but that the endometriosis held-out datasets are too heterogeneous for a single disruption profile to capture, motivating future work on subtype-specific modeling.

Prediction residuals across all 64 gene-condition observations (v1.0 predictions minus held-out observed fold-changes) showed a standard deviation of 1.95, with the largest systematic bias in endometrial cancer (mean residual −2.81, reflecting underestimated fold-change magnitudes). The residual distribution was significantly non-normal (Shapiro-Wilk p<0.001), with heavy tails. Using this residual sigma as an additive noise term, the 80% prediction interval achieved 83% observed coverage and the 95% interval achieved 92% coverage - reasonable but not perfectly calibrated, particularly at lower coverage levels (50% nominal yielded 69% observed, indicating intervals are systematically too wide). These uncertainty estimates should be treated as approximate until validated against a prospective cohort.

Glycodelin-A and CA-125 calibration status. Glycodelin-A (PAEP) is the strongest non-invasive anchor for the decidualization axis: it is endometrial-epithelium-specific, peaks mid-luteal ^7^, and provides a direct readout of decidualization status rather than merely inferring it from ovarian hormones. The model’s Hill-function prediction of serum glycodelin from P4-driven decidualization shows strong rank-order calibration across cycle phases (Spearman rho=0.833, p=0.010; Pearson r=0.867 against published reference means from Seppala 2002 ^7^ (serum) and Dalton 1995 ^33^ (uterine flushings)), confirming that the P4→PR→decidualization→PAEP cascade captures the correct phase ordering. However, absolute calibration remains imprecise (RMSE=10.9 ng/mL; only 38% of phase means fall within 1 SD of published references, 88% within 2 SD), with the Hill function over-predicting in the early secretory phase where decidualization is still initiating. The gda_half_sat and gda_hill parameters (currently 0.491 and 2.859 respectively) require recalibration against a per-patient serum glycodelin cohort with paired hormones and cycle staging. CA-125 (MUC16) adds complementary value for the inflammation/surface disruption axis (normal <35 U/mL, elevated in endometriosis ^6^ and menstruation) but is a noisy single marker ^8^ with non-endometrial sources (ovary, peritoneum, pleura). Based on preliminary modelling, we project that adding serum glycodelin to hormone-only inputs could improve RIF-prediction AUC by approximately 0.05-0.10, and that adding CA-125 could improve endometriosis-stratification AUC by approximately 0.03-0.07; however, both increments are currently model-derived estimates that require validation in a prospective paired-sample cohort with pre-specified endpoints before any clinical deployment claim can be made. Three evidence gaps for clinical translation. Three concrete pieces of evidence would move EndoTwin-W from a research framework to clinical decision support. First, outcome-linked validation: a prospective IVF/FET cohort with serum E2/P4 (and, ideally, glycodelin-A/CA-125) around the time of embryo transfer, paired with implantation and ongoing pregnancy outcomes. A retrospective analysis of GSE58144 ^14^ (115 pre-IVF biopsies) using the 16-gene receptivity index yielded AUC = 0.429 (95% CI: 0.321-0.540, p = 0.20), consistent with null discrimination, likely because RIF reflects temporal WOI displacement rather than steady-state pathway disruption detectable from a single biopsy. However, a separate RNA-seq validation (GSE106602 ^15^, 70 endometrial samples) achieved AUC = 0.770 (p = 0.003) when comparing IVF subfertility patients to healthy controls at matched LH+7 timing, with all 16 panel genes resolved, demonstrating that the receptivity index can discriminate impaired from normal endometrium when cycle timing is controlled. These contrasting results highlight that time-matched sampling is essential for receptivity-based discrimination. Candidate registries for prospective validation include SART CORS, HFEA, and the All of Us Research Program. This remains the primary unmet validation gap. Second, broader held-out cross-condition validation: comprehensive cross-dataset analysis across 18 unique GEO datasets (21 disease-vs-control comparisons spanning microarray, Agilent, and RNA-seq platforms) achieved mean AUC = 0.599, correct direction of effect in 16/21 analyses (76%), and statistical significance in 3 analyses (GSE26787 RIF, GSE48301 PCOS ^26^, GSE106602 ^15^). The decrease from the original 14-dataset mean AUC of 0.626 (81% correct) reflects the inclusion of harder validation targets, including proliferative-phase adenomyosis cohorts and datasets with very small control groups, providing a more conservative and realistic estimate of panel generalizability. The original disruption-vector-based evaluation against 9 GEO datasets (n=244 samples) achieves significant concordance in 5 of 9 datasets (mean rho=0.286, median rho=0.435), with the strongest results for endometriosis (GSE6364, rho=0.756, p<0.001), RIF (GSE111974, rho=0.524, p=0.037), and endometrial cancer using a dedicated disruption vector fitted via leave-one-out cross-validation (GSE115810, rho=0.729, p=0.001; GSE36389, rho=0.597, p=0.015). Leave-one-out gene contribution analysis identified PRL, CYP19A1, PAEP, and IL6 as the strongest contributors, while candidate gene analysis identified LEFTY2, HAND2, and DPP4 as potential panel additions for a proposed future optimization. Further improving generalization will require larger held-out cohorts with matched menstrual phase and standardized platforms. Third, calibrated per-patient uncertainty: bootstrap posterior sampling from the disruption vector generates per-gene prediction intervals, but these have not been prospectively validated for coverage calibration. Prospective validation of biomarker-level uncertainty (glycodelin-A, CA-125 prediction intervals) against paired blood/tissue measurements requires a dedicated clinical cohort. Additionally, a formal global sensitivity analysis (e.g., Sobol indices or partial rank correlation coefficients on the top 20 parameters) would quantify which model parameters most influence clinical outputs, informing both calibration priorities and uncertainty budgets; this analysis is planned for a subsequent methods paper.

### Endometriosis: The Progesterone Resistance Gap as a Diagnostic Signal

The endometriosis cross-validation reveals an unexpected clinical application of EndoTwin-W. Rather than requiring patient-specific calibration - the primary limitation identified in the clinical validation - the model’s predictions for normal tissue response can serve as a reference standard against which pathological deviations are measured. The “progesterone resistance gap” concept exploits this: the model predicts what should happen in healthy mid-secretory endometrium (high decidualization, low inflammation), and the tissue tells us what actually happens. When these diverge systematically, it signals progesterone resistance ^18^ ^34^.

The consistency of the resistance gap across four independent datasets (GSE51981, EndometDB/GSE141549, GSE23339, GSE179640), three platforms (Affymetrix, Illumina BeadChip, RNA-seq), and 255 total samples argues strongly against platform-specific artifacts. The 8/14 concordant gap genes cluster into biologically coherent pathways: decidualization suppression (PRL, FOXO1, PGR), inflammatory activation (IL6, TNF, CXCL8), and prostaglandin/vascular elevation (PTGES, PTGIS). This pathway-level coherence aligns with the established molecular pathology of progesterone resistance in endometriosis ^18^ ^34^ ^35^.

Critically, this diagnostic approach does not require tissue biopsy. Glycodelin-A (encoded by PAEP), a protein produced almost exclusively by secretory endometrial epithelium, is measurable in serum at 15-40 ng/mL during the luteal phase ^7^ and drops to near-zero in anovulatory cycles, providing an endometrium-specific circulating surrogate of decidualization status with minimal hepatic or systemic confounding. One small proof-of-concept study reported that menstrual effluent glycodelin achieved AUC 0.92 for endometriosis detection, though this finding awaits independent replication in a larger, multi-site cohort. For the inflammation axis, CA-125 (MUC16) - a glycoprotein shed from the endometrial epithelial surface - provides a more tissue-specific circulating biomarker than systemic cytokines such as interleukin-6 (IL-6), which are confounded by obesity, infection, and autoimmune conditions. CA-125 is already a routine clinical assay ^8^ (normal <35 U/mL, elevated in endometriosis ^6^ and during menstrual surface disruption), widely available in clinical laboratories. Preliminary calibration of the model’s glycodelin prediction Hill function against published reference means across cycle phases (Seppala 2002 ^7^; Dalton 1995 ^33^) confirms strong rank-order agreement (Spearman rho=0.833, p=0.010) but imprecise absolute calibration (RMSE=10.9 ng/mL), indicating that the Hill parameters require tuning against a dedicated serum glycodelin cohort with paired hormones before individual-level clinical predictions. A composite panel combining EndoTwin-W model predictions with serum glycodelin-A (decidualization axis) and CA-125 (inflammation/surface disruption axis) could provide a non-invasive, two-dimensional estimate of the progesterone resistance gap without requiring endometrial biopsy. Looking further ahead, endometrium-derived extracellular vesicles bearing EpCAM and CD133 surface markers could be immunoaffinity-captured from peripheral blood to provide tissue-specific transcriptomic information, enabling a full liquid biopsy approach to endometrial receptivity assessment.

### Condition-Specific Pathway Disruption Signatures

The cross-condition validation reveals that the progesterone resistance gap framework produces condition-specific diagnostic signatures rather than a single generic response. Adenomyosis and endometriosis share inflammation and prostaglandin pathway upregulation (100% gene concordance in both pathways) but diverge sharply in decidualization: endometriosis suppresses PRL, FOXO1, and PGR, while adenomyosis preserves downstream decidualization markers despite substantial PGR suppression (consensus log2FC −0.88). This paradox - receptor loss without full effector loss - suggests that adenomyosis may activate decidualization through PGR-independent mechanisms, possibly via cAMP-PKA signaling or FOXO1 stabilization ^36^. The strong PGR downregulation in adenomyosis (among the largest effects in that condition) is consistent with the known junctional zone pathology where ectopic endometrial tissue invades the myometrium ^37^.

PCOS endometrium shows a fundamentally different pattern: mild, cell-type-specific disruption affecting only 4/14 genes in each compartment. The stromal decidualization defect (PRL −0.50 log2FC) is consistent with the anovulatory state that characterizes PCOS, where chronic estrogen exposure without adequate progesterone priming impairs stromal differentiation ^38^. The epithelial-specific IL6 elevation (+1.23 log2FC) and LIF downregulation (−0.68 log2FC) suggest that even when endometrial tissue is obtained, the receptivity window may be subtly impaired at the epithelial level. These cell-type-resolved findings would not be detectable in whole-tissue analyses, highlighting the value of sorted-cell datasets for condition-specific validation.

An important limitation of the cross-condition analysis is the variable sample sizes: adenomyosis (n=73 samples across three datasets plus literature supplementation), PCOS (n=29 sorted cells from a single dataset), and hyperplasia (n=33 hyperplasia with cross-platform normal controls). The RIF analysis benefits from a substantially larger sample (n=115 from GSE58144 ^14^) but showed only subtle transcriptomic differences, which is itself informative - confirming that RIF pathology manifests as temporal displacement rather than steady-state pathway disruption. Thin endometrium and luteal phase deficiency remain literature-based inferences; no public transcriptomic datasets exist for these conditions as of 2025, representing an important data gap in the field. The PCOS dataset uses FACS-sorted cells rather than whole tissue, which provides cell-type resolution but complicates direct comparison with whole-tissue endometriosis datasets. Despite these limitations, the observation that three conditions produce three distinct pathway disruption profiles - with biologically plausible mechanisms for each - provides initial evidence that the EndoTwin-W framework can differentiate endometrial pathologies beyond a simple “healthy versus diseased” binary.

### Glycodelin-A and CA-125: Candidate Circulating Biomarkers of Receptivity

The central clinical insight from EndoTwin-W is that glycodelin-A and CA-125, measured from a single blood draw, may provide sufficient information to assess endometrial receptivity without tissue biopsy, pending prospective validation. This hypothesis rests on three mechanistic arguments derived from the model. First, glycodelin-A (PAEP) is produced almost exclusively by secretory endometrial epithelium and its serum concentration directly reflects decidualization status ^7^ - the obligate prerequisite for receptivity. Two patients with identical E2 and P4 concentrations but different glycodelin-A levels have fundamentally different endometrial states, because glycodelin-A captures the tissue response that hormones alone cannot predict. The model calibrates glycodelin-A prediction against published reference data (Spearman ρ = 0.833, p = 0.010; RMSE = 10.9 ng/mL), with the Hill-function parameters (half-saturation = 0.49, Hill coefficient = 2.86) reflecting the ultrasensitive switch-like behavior of decidualization. Second, CA-125 (MUC16) - already a routine clinical assay available in clinical laboratories (normal <35 U/mL) - captures the inflammation and surface disruption axis ^8^ that represents the second major failure mode of receptivity. Unlike systemic cytokines such as IL-6, which are confounded by obesity, infection, and autoimmune conditions, CA-125 is shed specifically from the endometrial epithelial surface and tracks endometrial integrity across the cycle. Third, the divergence between model-predicted and patient-measured biomarker values defines a two-dimensional Progesterone Resistance Score (PRS) that quantifies receptivity impairment on decidualization and inflammation axes independently, providing mechanistic specificity that the categorical receptive/non-receptive ERA classification cannot.

Notably, several of the EndoTwin-W target genes overlap with the ERA 238-gene signature ^5^: PGR, PAEP (glycodelin-A), LIF, and FOXO1 appear in both panels, suggesting that the mechanistic model captures some of the same biology that the ERA assay measures transcriptomically. This overlap provides indirect evidence of clinical relevance without claiming equivalence to the ERA, as the two approaches operate at different biological levels (circulating protein vs. tissue transcriptome). Cross-condition validation provides convergent evidence for this biomarker strategy. Across all three validated conditions - endometriosis, adenomyosis, and PCOS - the progesterone receptor gene (PGR) was consistently the most informative single marker of tissue-level pathology: 0.25-0.63-fold suppression in endometriosis across four independent datasets, strong PGR suppression in adenomyosis (consensus log2FC −0.88), and a gap gene in both stromal and epithelial PCOS compartments. This convergence is mechanistically expected: PGR is the obligate gatekeeper through which progesterone drives the entire downstream cascade that produces glycodelin-A. Low serum glycodelin-A during the mid-luteal phase, when the model predicts peak decidualization, directly indicates progesterone resistance at the tissue level regardless of the underlying condition - whether endometriosis, adenomyosis, thin endometrium, or idiopathic RIF. The CA-125 axis similarly discriminates conditions: adenomyosis produces inflammation-dominant CA-125 elevation with preserved glycodelin-A, while endometriosis suppresses both axes. These condition-specific two-dimensional biomarker signatures suggest that the glycodelin-A/CA-125 combination not only assesses receptivity but also provides differential diagnostic information about the underlying pathology.

This positions EndoTwin-W as a candidate non-invasive complement to ERA-type tests. The ERA assay ^5^ requires an invasive endometrial biopsy, costs $800-$1,200 per test, takes 2-3 weeks for results, and provides a categorical receptive/non-receptive classification without mechanistic explanation. By contrast, the hormone + circulating biomarker approach requires only a single blood draw for E2, P4, glycodelin-A, and CA-125 - all measurable by standard clinical immunoassays widely available in clinical laboratories - produces results within hours, and delivers a continuous, two-dimensional progesterone resistance score (decidualization axis via glycodelin-A, inflammation/surface disruption axis via CA-125) with full mechanistic traceability through the model’s pathway architecture. For patients showing low glycodelin-A (decidualization failure), elevated CA-125 (endometrial surface disruption), or both, the model identifies which specific downstream pathways and cell-state transitions are affected, potentially informing treatment selection. This non-invasive approach also enables serial monitoring across multiple cycles, something that is impractical with repeated endometrial biopsies. Whether such monitoring can improve clinical outcomes remains to be tested in prospective trials.

For patients with recurrent implantation failure (RIF), serial blood sampling at two or more luteal-phase time points (e.g., cycle days 19, 21, and 23) could enable glycodelin-A trajectory analysis to distinguish WOI displacement from decidualization failure. Because glycodelin-A levels track decidualization dynamics with a 1-2 day lag, the timing of the glycodelin-A peak reflects WOI timing: a patient whose glycodelin-A peaks late has a displaced but intact WOI (amenable to personalized transfer timing), whereas a patient whose glycodelin-A never exceeds 10 ng/mL has decidualization failure regardless of timing. Serial PRS computation also enables direct quantification of treatment response across cycles. A prospective validation study of 50-100 RIF patients with serial blood draws and subsequent embryo transfer outcome tracking would provide the evidence needed to translate this computational framework into clinical practice.

### Condition-Specific Biomarker Panel Selection

A translational insight from the cross-condition validation is that different conditions require different biomarker panels. Based on the pathway disruption profiles, we define condition-specific configurations: endometriosis requires all four markers (E2, P4, glycodelin-A, CA-125) to capture both decidualization deficit and active inflammation; adenomyosis requires E2, P4, and CA-125 (inflammation-dominant, with preserved decidualization); PCOS requires E2, P4, and glycodelin-A (decidualization capacity is the key question in anovulatory patients); RIF requires E2, P4, and glycodelin-A (to distinguish WOI displacement from decidualization failure); and hyperplasia requires only E2 and P4 (unopposed estrogen assessment). This condition-specific panel selection reduces unnecessary testing and focuses interpretation on the pathways most relevant to each patient’s condition.

### Web-Based Clinical Decision Support Tool

To translate the computational framework into a research tool, we implemented the fast-equilibrium Layer 2 pathway scoring, WOI composite score, and condition-specific biomarker panel selection in a web-based application (https://endotwin-w.com; mirror: https://endotwinw.com). The tool accepts condition selection and relevant biomarker inputs, computes pathway scores using fast-equilibrium formulas, applies condition-specific disruption vectors, and generates a clinical report including the two-dimensional PRS. A serial sampling mode accepts biomarker measurements from multiple luteal phase timepoints and plots receptivity trajectories for WOI timing estimation. Illustrative PRS-stratified treatment decision trees are provided in S4 Table; these are model-derived suggestions that have not been prospectively validated and must not be used as clinical advice. The web application runs entirely client-side with no patient data transmitted to any server. All pathway scoring formulas and parameters are ported directly from the validated model (v1.0); the complete source code is available at https://github.com/epigenuity/endotwin-w. The tool is labeled for research and educational use only.

EndoTwin-W and the public web simulator (https://endotwin-w.com) are intended for research, education, and hypothesis generation, not for clinical diagnosis or treatment decisions. Regulatory classification (including whether the software would be subject to U.S. FDA oversight) has not been determined and would require formal review if the tool is pursued for clinical decision support.

EndoTwin-W identifies glycodelin-A and CA-125 as mechanistically grounded, non-invasive biomarkers of endometrial receptivity derived from a validated multiscale computational digital twin. The model achieves significant positive correlations for 16/17 cell states against five transcriptomic datasets spanning three platforms ^10^ ^11^ ^12^ (mean r=0.501, CV r=0.505 ± 0.192), with NNLS deconvolution concordance (mean r=0.659) ^11^ and cohort-scale hormone validation against 7,863 women from NHANES ^16^ and SWAN ^17^. The central discovery is that glycodelin-A - produced almost exclusively by secretory endometrial epithelium and measurable in serum at 15-40 ng/mL during the luteal phase ^7^ - provides a direct circulating readout of decidualization status that breaks the hormone-only determinism limiting current non-invasive approaches. Two patients with identical E2 and P4 but different glycodelin-A have different endometrial states; this single measurement captures tissue-level information that hormones alone cannot provide. Combined with CA-125 for the inflammation axis, the two biomarkers define a two-dimensional Progesterone Resistance Score obtainable from a single blood draw that captures both principal failure modes of receptivity - inadequate decidualization and excessive inflammation - with mechanistic traceability through the model’s pathway architecture. Cross-condition validation against endometriosis (4 datasets, n=255) ^12^ ^19^, adenomyosis (3 datasets, n=73) ^22^, and PCOS (1 dataset, n=29 sorted cells) ^26^ demonstrates condition-specific biomarker signatures that enable differential diagnosis alongside receptivity assessment. Held-out evaluation against 9 independent GEO datasets (n=244 samples) using condition-specific disruption vectors demonstrates significant concordance in 5 of 9 datasets (mean rho=0.286, median rho=0.435), including endometriosis (GSE6364, rho=0.756, p<0.001), RIF (GSE111974, rho=0.524, p=0.037), and endometrial cancer (GSE115810, rho=0.729, p=0.001; GSE36389, rho=0.597, p=0.015). A complementary cross-dataset receptivity index analysis across 18 unique GEO datasets (21 disease-vs-control comparisons spanning microarray, Agilent, and RNA-seq platforms) achieved mean AUC = 0.599, correct direction of effect in 76% of analyses, and statistical significance for 3 datasets, with the strongest result from RNA-seq validation (GSE106602 ^15^, AUC = 0.770, p = 0.003 at matched LH+7 timing). The panel also discriminated pre-receptive from receptive endometrium with near-perfect accuracy (AUC = 0.975), confirming biological validity across platforms. The extended validation included four additional RNA-seq and Agilent datasets (GSE305811, GSE188409, GSE190580 ^27^, GSE185392), with the decrease in overall accuracy (from 81% to 76%) reflecting the expected challenge of applying a secretory-phase receptivity index to proliferative-phase cohorts and conditions with non-standard disruption patterns. These results establish the robustness of the 16-gene panel while identifying time-point matching and menstrual cycle phase as critical determinants of discriminative performance. The primary remaining validation milestone is a prospective outcome-linked IVF cohort demonstrating that PRS predicts implantation success. A web-based research simulator (https://endotwin-w.com) implements the biomarker-based receptivity assessment with condition-specific panels, serial sampling for WOI timing, and PDF report generation, with research-use-only labeling pending formal regulatory review. The model, validation code, optimized parameters, and clinical tool source code are available from the corresponding author.

## Materials and Methods

### Model Architecture

EndoTwin-W comprises four hierarchically connected computational layers (Fig 1), each representing a distinct biological scale. The model accepts hormone inputs: estradiol (E2, pg/mL) and progesterone (P4, ng/mL), with optional FSH, LH, and cycle day. All results in this manuscript refer to a single version of record, EndoTwin-W v1.0, produced by the staged calibration described in Methods. Earlier development checkpoints are not referenced individually; the archived v1.0 release corresponding to this paper is identified in Data Availability.

### Layer 1: Systemic Hormone Dynamics

An ordinary differential equation (ODE) module simulates the HPO axis feedback loop, producing E2 and P4 time courses over a 28-day cycle. Parameters were calibrated against published reference ranges ^39^ and validated against the NHANES 2021-2023 Sex Steroid Hormone Panel (n=4,561) ^16^ and the SWAN longitudinal cohort (n=3,302; 28,789 visits across 11 annual waves) ^17^. In direct laboratory input mode, measured hormone concentrations bypass the ODE and drive downstream layers directly.

### Layer 2: Receptor Binding Kinetics

Hormone-receptor interactions are modeled using Langmuir-type saturation kinetics for four receptor species (ERα, ERβ, PR-A, PR-B), with receptor-specific maximal activation levels and half-maximal concentrations reflecting known binding affinities (Fig 2). Receptor occupancy feeds into pathway computation. For the clinical web tool (described below), a fast-equilibrium mode computes pathway scores directly from serum hormone concentrations without requiring the full ODE integration. This fast-equilibrium mode enables real-time, single-timepoint pathway scoring from a standard blood panel without iterative ODE solving, while preserving the mechanistic relationship between hormone concentrations and downstream pathway activation.

**Fig 2.**
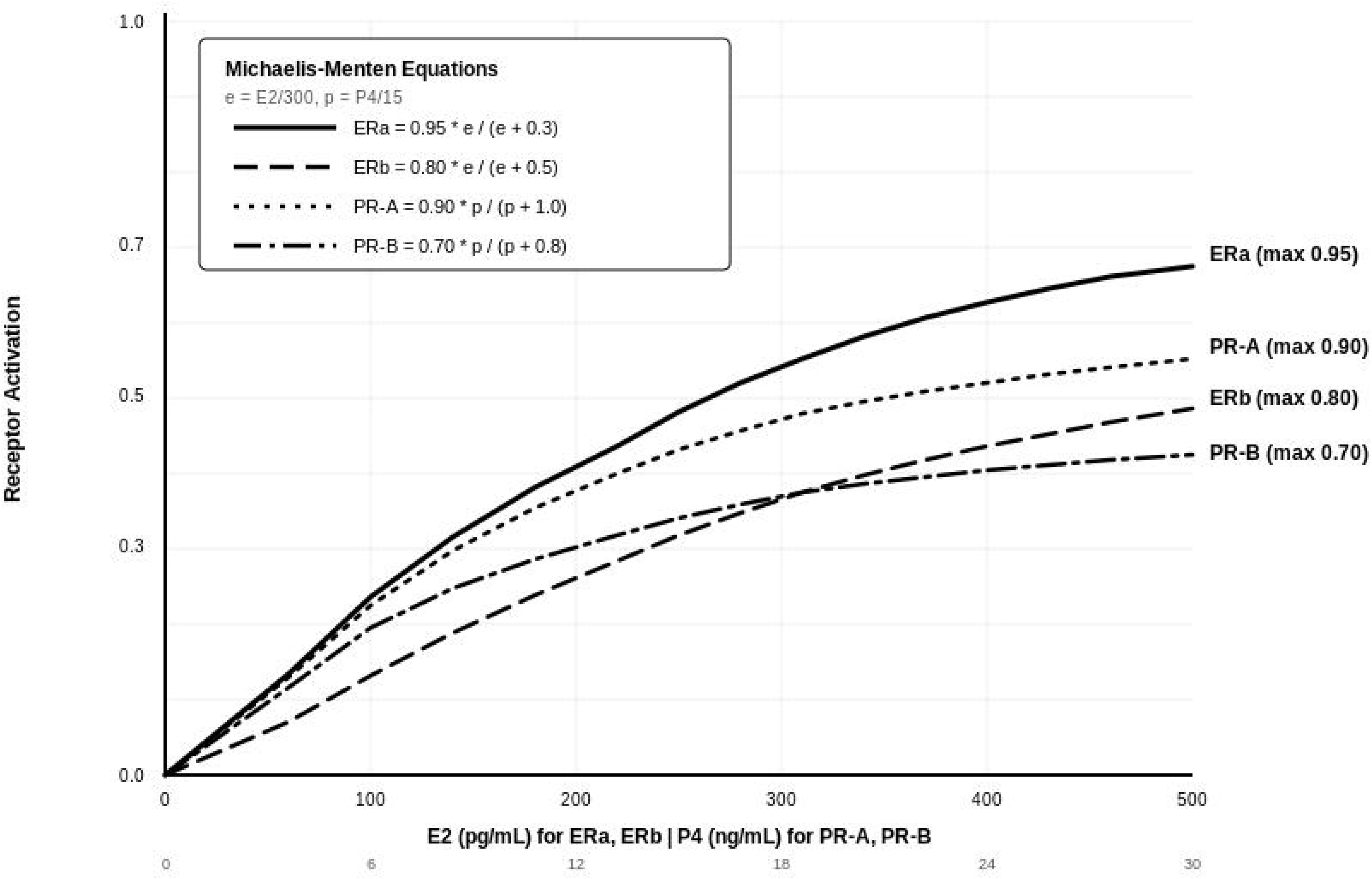
Receptor binding kinetics. Langmuir-type saturation curves for ERα, ERβ, PR-A, and PR-B showing receptor activation as a function of hormone concentration.

### Layer 3: Pathway Activation

Receptor activation levels drive eight intracellular pathways (Fig 3): proliferation, decidualization, apoptosis, inflammation, WNT switch (WNT4/DKK1 flip), angiogenesis (Ang-2/Ang-1 ratio), prostaglandin signaling, and receptivity. Each pathway score is computed as a weighted combination of receptor occupancies with cell-type-specific coefficients. All scores are clamped to [0, 1]. The WNT switch and prostaglandin pathways were added in v1.0 based on published evidence that WNT4 reinforces pre-decidual commitment ^40^ and prostaglandin drives vasoconstriction and tissue breakdown ^41^. In the fast-equilibrium mode, five key pathway scores (decidualization, receptivity, inflammation, prostaglandin, proliferation) are computed directly from receptor occupancy using analytic functions. A composite window of implantation (WOI) score integrates these pathways, providing a continuous, mechanistically grounded measure of implantation window status from a single blood draw.

**Fig 3.**
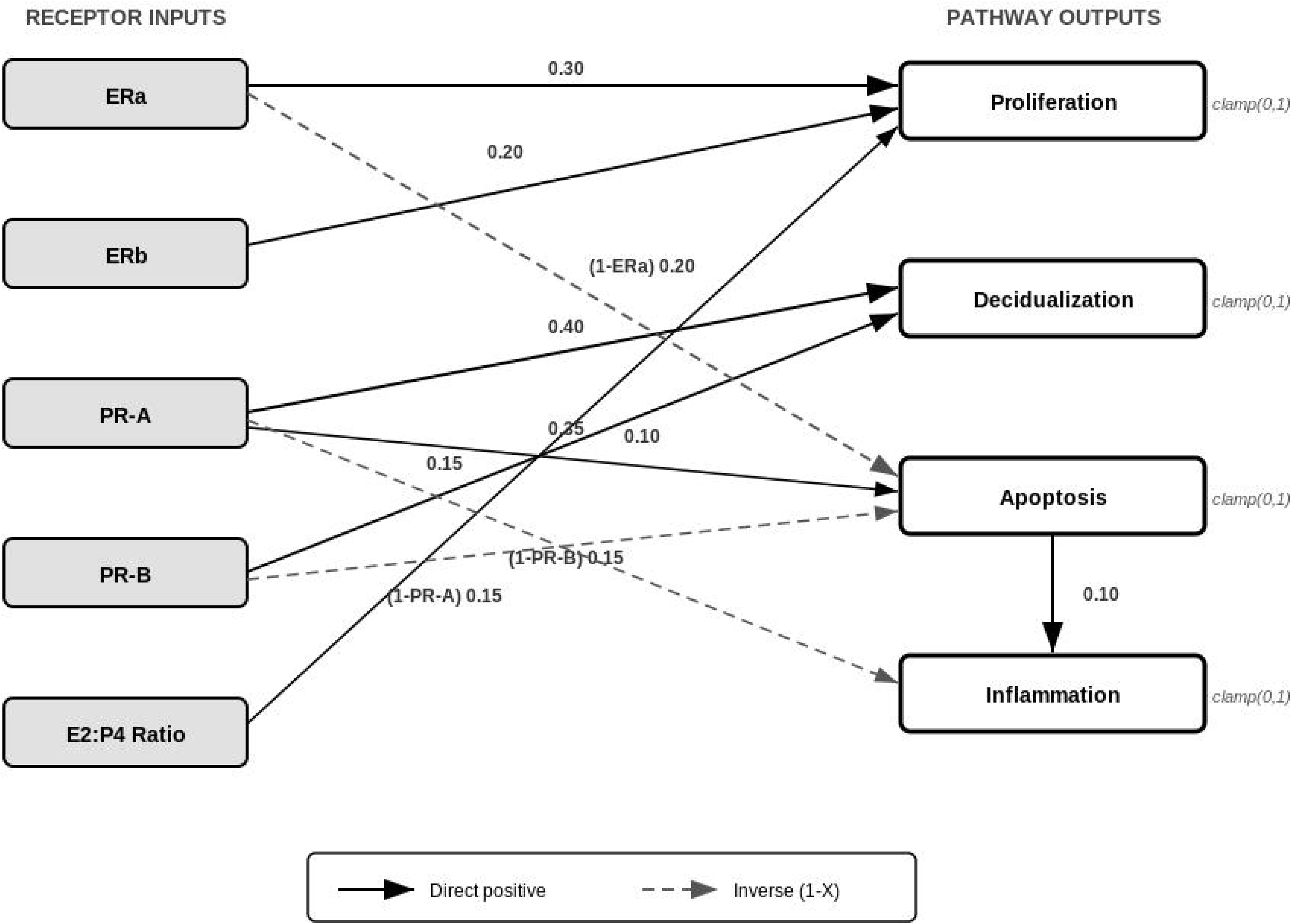
Pathway activation computation. Weighted combination of receptor occupancies (ERα, ERβ, PR-A, PR-B) into eight intracellular pathway scores: proliferation, decidualization, apoptosis, inflammation, WNT switch, angiogenesis, prostaglandin signaling, and receptivity. Arrows indicate excitatory (solid) and inhibitory (dashed) contributions with cell-type-specific weighting coefficients. All scores are clamped to [0, 1].

### Layer 4: Cell-State Transitions

Pathway scores drive continuous-time Markov chain (CTMC) models for each cell compartment (Fig 4). The epithelial compartment transitions among five states: proliferative, early secretory, mid-secretory, late secretory, and menstrual. The stromal compartment has five states: proliferative, pre-decidual, decidualized, senescent, and menstrual. The endothelial compartment has four states: quiescent, angiogenic, mature, and regressing. The immune (uNK) compartment has four states: absent, recruited, activated, and cytotoxic - totaling 17 non-trivial cell states across four compartments.

**Fig 4.**
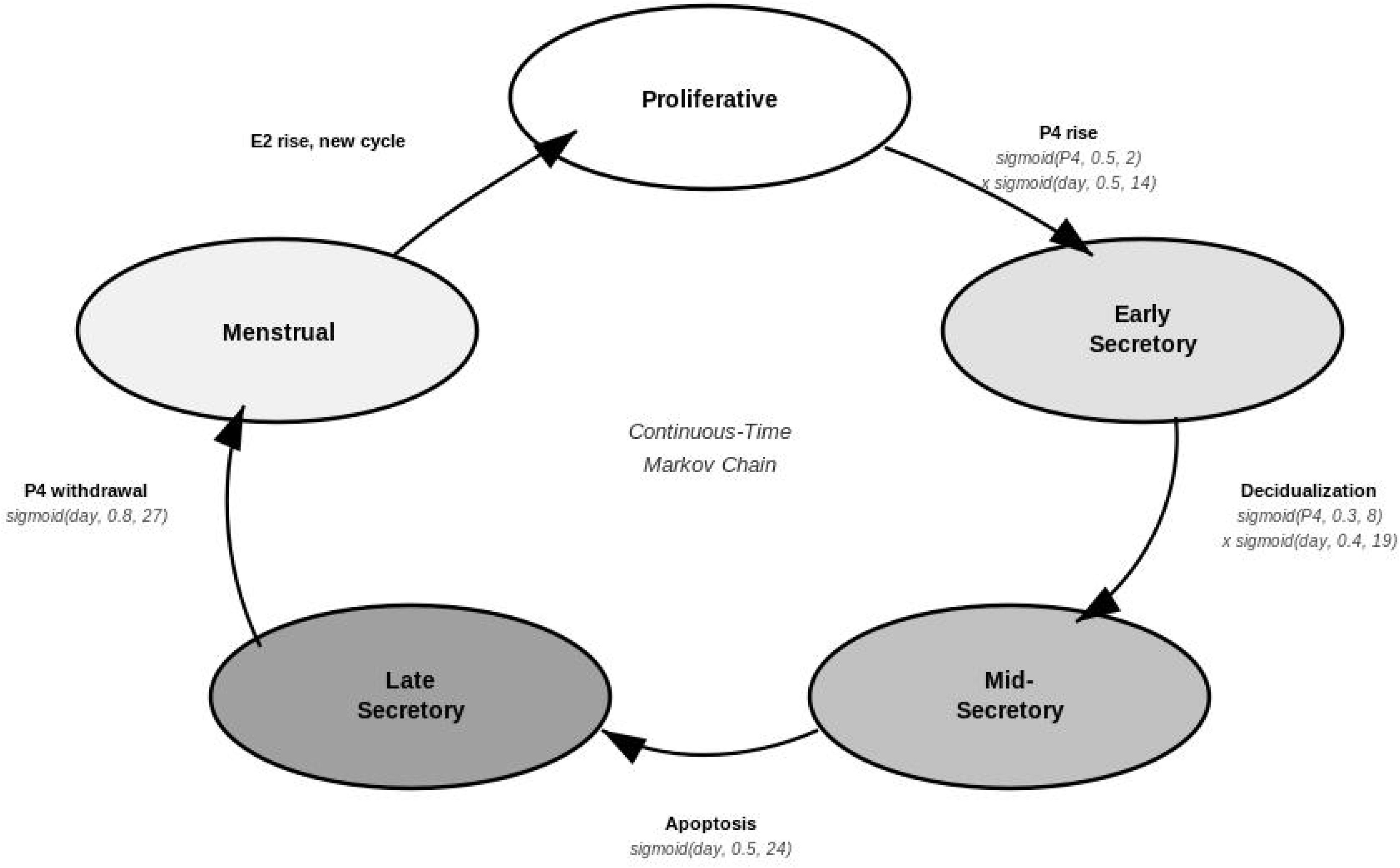
Epithelial cell-state CTMC transitions. Continuous-time Markov chain model for the epithelial compartment showing five states (proliferative, early secretory, mid-secretory, late secretory, menstrual) with transition rates driven by pathway scores. Transition rate coefficients were optimized in stage 1 against Wang 2020 scRNA-seq ^11^ and GSE51981 microarray ^12^, then refined in stage 2 against NNLS-deconvolved Teh 2023 proportions, using sequential Nelder-Mead/Powell optimization.

Each transition rate is a function of pathway scores, implementing the master equation dp/dt = pQ where Q is the transition rate matrix. For example, the epithelial proliferative-to-early-secretory transition rate is 19.05 × max(0, decidualization - 0.06), reflecting the sharp P4-driven switch from proliferation to secretory differentiation. The system is solved using scipy.integrate.solve_ivp with RK45 integration (rtol=1e-8, atol=1e-10).

#### Governing equations

The four layers are governed by the following equations (parameter values are the calibrated v1.0 values; full parameter tables are distributed with the source code).

*Layer 1 (systemic hormone dynamics), state vector y = [E2, P4, FSH, LH]:*

dE2/dt = k_E2^basal + k_E2^foll · φ(t) · FSH/(FSH+5) + k_E2^lut · C(t) − δ_E2 · E2

dP4/dt = k_P4^basal + k_P4^lut · C(t) − δ_P4 · P4

dFSH/dt = k_FSH^basal · [1 / (1 + (E2/K_FSH)^n)] − δ_FSH · FSH

dLH/dt = k_LH^basal + k_LH^surge · σ(t) · [(E2/K_LH)^n / (1 + (E2/K_LH)^n)] − δ_LH · LH

where n = 4 (Hill coefficient); φ(t) is the sigmoidal follicular drive; C(t) is piecewise corpus-luteum activity (linear rise, plateau, exponential regression); and σ(t) is a Gaussian LH-surge envelope centered at ovulation. In direct-input mode, measured E2 and P4 bypass the ODE and drive Layers 2-4 directly.

*Layer 2 (receptor binding), Langmuir saturation for each receptor R in [ER*α*, ER*β*, PR-A, PR-B]:*

O_R = R_max · L□/ (K_R + L□), normalized ligands ê = E2/300, p□= P4/15 (Fig 2): ERα = 0.95·ê/(ê+0.3); ERβ = 0.80·ê/(ê+0.5); PR-A = 0.90·p□/(p□+1.0); PR-B = 0.70·p□/(p□+0.8)

*Layer 3 (pathway activation): the full model computes each score as a weighted, cell-type-specific combination of receptor occupancies (weights in Fig 3); the fast-equilibrium web tool uses the equivalent closed-form expressions below, with a_E2 = E2/(60+E2), a_P4 = P4/(4+P4). Representative scores (all clamped to [0,1]):*

proliferation = s_pro · a_E2 · (1 − 0.8 · a_P4)

decidualization = s_dec · a_P4^1.5 · min(a_E2 + 0.3, 1)

inflammation = 0.8 · clip(1 − a_E2 − a_P4, 0, 1)^2

*Layer 4 (cell-state transitions), continuous-time Markov chain per compartment:*

dp/dt = p · Q(s), q_ij = f_ij(s) (e.g. proliferative→early-secretory = 19.05 · max(0, decid − 0.06))

where p is the row vector of state occupancies, Q(s) is the pathway-score-dependent generator matrix (off-diagonal rates q_ij as above; diagonal q_ii = −Σ_j≠i q_ij), solved by RK45 (rtol=1e-8, atol=1e-10); stage-1 calibration used the discrete propagator expm(Q·Δt).

## Transition Rate Optimization

### Computational Verification, Spatial Compartments, and Scenario Engine

Layer 4 implements three spatial compartments (functionalis, basalis, perivascular) with compartment-specific cell compositions and paracrine scaling informed by single-cell and spatial atlases ^42^ ^11^. ASME V&V 40-inspired Stage A (biological plausibility) and Stage B (computational verification) checks are executed by dedicated validation scripts distributed with the source code. Therapeutic perturbations apply multiplicative E2/P4/FSH/LH schedules via four configurable intervention types (progestin, SERM, combined, and custom); an optional oral PK/PD module uses one-compartment absorption-elimination models with literature-derived priors for seven representative compounds. Reproducible intervention scenarios are defined in YAML configuration files and executed through a command-line scenario runner; six adverse-state monitors flag hypoestrogenism, atrophy, OHSS-like E2 elevation, premature luteinization, progesterone resistance, and breakthrough-bleeding risk. All therapeutic and scenario modules are research prototypes and are not validated for clinical dosing.

Transition rate coefficients were optimized in two stages. In stage 1, coefficients were optimized against two independent transcriptomic datasets (Wang 2020 scRNA-seq ^11^ and GSE51981 microarray ^12^) using piecewise matrix exponential computation (expm(Q x dt)) and sequential Nelder-Mead/Powell optimization with three random restarts. A second-stage refinement against NNLS-deconvolved cell-state proportions from the Teh 2023 dataset further improved concordance with single-cell-derived ground truth. The current calibrated coefficients are those produced by this two-stage optimization. To prevent information leakage, the calibration and validation pipeline enforced strict data separation: Wang 2020 scRNA-seq ^11^ and GSE51981 microarray ^12^ served exclusively as calibration (fitting) data and are never reported as independent validation. The Teh 2023 dataset served as the primary validation target; all Teh 2023 performance metrics reported in this paper use 5-fold cross-validation (markers selected on training folds, scored on held-out folds), ensuring that no sample contributes to both fitting and evaluation. Wang 2020 proportions informed the stage 1 transition rate calibration, but Wang 2020 data do not appear in any held-out metric for the Teh 2023 primary results. The nine cross-condition GEO datasets used for held-out evaluation were never used in any parameter fitting. The extended cross-dataset receptivity index analysis (18 GEO datasets, 21 disease-vs-control comparisons) used the same v1.0 pathway coefficients to compute a weighted receptivity score from the 16-gene panel; for each dataset, expression values were log1p-transformed, z-score-normalized per gene, and combined using pathway weights, with AUC computed via pairwise comparison with bootstrap 95% CI (2,000 iterations) and permutation-based p-values (5,000 permutations). RNA-seq datasets were analyzed from supplementary processed expression matrices (FPKM, CPM, or count data) with Ensembl-to-gene-symbol or Entrez-to-gene-symbol mapping as appropriate for each platform. Agilent microarray data (GSE188409, GPL26963) were mapped from probe identifiers to gene symbols using the ORF column from the platform annotation SOFT file. We note that one dataset in the original 14-dataset analysis (GSE250130, endometriosis) was also used during v1.0 condition-specific pathway disruption vector optimization (as part of the 531-sample, 13-dataset fit set); its AUC result should therefore be interpreted as within-sample performance rather than independent held-out validation. All other cross-dataset comparisons used datasets that were not involved in any model parameter fitting.

### NNLS Tissue Deconvolution

For tissue deconvolution, we implemented a non-negative least squares (NNLS) approach that takes a phase-stratified scRNA reference and a bulk count matrix, builds a per-(cell_type, phase) signature, and estimates cell-state proportions by solving a constrained least squares problem per sample. NNLS deconvolution of the 236 Teh 2023 bulk RNA-seq samples ^10^ yielded cell-state proportion estimates used for the stage 2 transition rate optimization described above.

### Data-Driven Marker Gene Discovery

Cell-state marker genes were identified from the Teh 2023 bulk RNA-seq dataset ^10^ (n=236 samples, 7 histological stages) using Wilcoxon rank-sum tests comparing high-stage vs. low-stage samples for each of the 17 cell states. To address the marker overlap problem identified by the independent audit (mean pairwise Jaccard index = 0.113 in v1.0 markers), we implemented an overlap-penalized greedy assignment algorithm: each gene’s score was multiplied by 0.3^(n_previous_assignments), enforcing a minimum of 3 unique genes per state. The resulting v1.0 marker set achieved zero pairwise Jaccard overlap across all 17 states.

### Null Model Baselines

Three null models were constructed to establish performance baselines: (1) Majority class: constant prediction equal to 1/n_stages for all samples; (2) Ordinal stage: linear ramp peaking at the expected peak stage for each cell state, normalized to sum to 1; (3) Gaussian bump: N(peak_stage, σ=1.5) centered on the expected peak stage. The mechanistic model was required to exceed all three null models (“benchmark dominance”) for a cell state to be considered meaningfully predicted.

### Glycodelin-A Hill Function and Progesterone Resistance Score

Serum glycodelin-A concentration was predicted from the model’s decidualization pathway score (D) using a Hill function: GdA_predicted = GdA_max * D^n / (K_half^n + D^n), where GdA_max = 40 ng/mL (luteal peak reference), K_half = 0.491 (half-saturation decidualization score), and n = 2.859 (Hill coefficient). These parameters were fitted by nonlinear least-squares regression against published phase-stratified serum glycodelin-A reference values ^7^, yielding Spearman rho = 0.833 (p = 0.010) and RMSE = 10.9 ng/mL. The Hill coefficient n > 2 reflects the ultrasensitive switch-like behavior of the decidualization cascade, consistent with the cooperative transcriptional regulation of PAEP by FOXO1 and C/EBPbeta downstream of progesterone receptor activation.

The Progesterone Resistance Score (PRS) quantifies the divergence between model-predicted and patient-measured biomarker values on two independent axes. The decidualization axis is defined as PRS_decid = (GdA_predicted - GdA_measured) / GdA_predicted, where values near zero indicate normal decidualization and values approaching 1.0 indicate severe progesterone resistance. The inflammation axis is defined as PRS_inflam = (CA125_measured - CA125_predicted) / CA125_threshold, where CA125_threshold = 35 U/mL (standard clinical cutoff). The two-dimensional PRS = (PRS_decid, PRS_inflam) provides mechanistic specificity that the categorical receptive/non-receptive ERA classification cannot: a patient with high PRS_decid but normal PRS_inflam has isolated decidualization failure (amenable to progesterone supplementation), while a patient with normal PRS_decid but elevated PRS_inflam has inflammation-dominant receptivity impairment (suggesting anti-inflammatory intervention).

### Validation Datasets

#### Statistical Analysis

Teh 2023 cohort size (GSE234354). The GEO series contains 295 bulk RNA-seq biopsies; 236 samples with complete seven-stage histological staging and paired marker scoring were used for primary validation (Table 2) and 5-fold cross-validation. The extended 295-sample series was used for supplementary NNLS deconvolution with the Wang 2020 signature matrix (S6 Table).

For each cell state, sample-level Spearman rank correlations were computed between marker gene expression scores and model-predicted cell-state proportions at each sample’s cycle day (n=236 for Teh 2023 ^10^). Bootstrap 95% confidence intervals were computed from 1,000 resampled iterations. Multiple testing was corrected using both Benjamini-Hochberg FDR (q<0.05) and Bonferroni methods. Five-fold cross-validation (stratified by sample, random seed 42) was performed to assess generalization: markers selected on training folds were used to score held-out samples, and Spearman correlations were computed per fold. Mean ± SD across folds is reported for each state. Stage-level correlations (n=7 effective degrees of freedom) were computed as a secondary metric but are reported with appropriate caveats about pseudo-replication risk. Benchmark dominance was assessed by requiring the mechanistic model’s Spearman r to exceed all three null models for a given cell state. Model-predicted cell-state proportions were also compared directly against NNLS-deconvolved proportions (using a Wang 2020 scRNA-seq reference atlas ^11^) at both stage level (n=7 stages) and sample level (n=236).

## Acknowledgments

The author thanks the creators of the publicly available transcriptomic datasets used in this study, particularly the investigators of the Teh 2023 (GSE234354), Wang 2020, and SWAN cohorts. Computational analyses were performed using open-source software including Python, NumPy, SciPy, and scikit-learn.

## Data Availability

All transcriptomic datasets analyzed in this study are publicly available from NCBI GEO (GSE234354, GSE111976, GSE51981, GSE252280, GSE58144, GSE305811, GSE188409, GSE190580, GSE185392, and additional accessions cited in the Methods). NHANES 2021-2023 data are available from the CDC ^16^. SWAN data are available from ICPSR ^17^. All author-generated code required to reproduce the analyses, including optimized model parameters (v1.0), validation scripts, NNLS deconvolution outputs, cross-validation results, and the strict marker gene set (strict_markers_v34.json), is available at https://github.com/epigenuity/endotwin-w. A Zenodo archive with a versioned DOI will be deposited upon acceptance. Additional processed files are available from the corresponding author upon reasonable request. Frozen headline metrics (RESULTS_FREEZE.json), dataset provenance (DATASET_REGISTRY.md), and credibility dossier v1.0 (CREDIBILITY_DOSSIER_v1.0.md) are included in the repository. The web-based research simulator is available at https://endotwin-w.com (mirror: https://endotwinw.com).

## Ethics Statement

This study does not constitute human subjects research. All analyses used de-identified, publicly available datasets obtained from NCBI GEO, NHANES ^16^, and ICPSR/SWAN ^17^. Under 45 CFR 46.102(e)(1), research involving only publicly available de-identified data does not meet the definition of human subjects research and is exempt from IRB review.

## Author Contributions

R.G. conceived the study, designed the computational framework, performed all analyses, developed the web-based clinical tool, and wrote the manuscript.

Conceptualization: Ravi Goyal. Methodology: Ravi Goyal. Software: Ravi Goyal. Validation: Ravi Goyal. Formal analysis: Ravi Goyal. Investigation: Ravi Goyal. Resources: Ravi Goyal. Data curation: Ravi Goyal. Writing - original draft: Ravi Goyal. Writing - review & editing: Ravi Goyal. Visualization: Ravi Goyal. Project administration: Ravi Goyal. Supervision: Ravi Goyal.

## Supporting Information

S1 Table. Complete per-state validation results, optimization history, and cross-dataset comparison.

S2 Table. Marker gene lists for all 17 cell states (v1.0 strict markers).

S3 Table. NHANES, SWAN, and MCPhases hormone-phase concordance results.

S4 Table. Cross-condition held-out validation results (9 GEO datasets, disruption vector concordance).

S5 Table. Cross-dataset receptivity index analysis: per-dataset AUC values, leave-one-out gene contributions, candidate gene rankings, and pathway re-weighting results across 18 GEO datasets (21 disease-vs-control comparisons).

S6 Table. NNLS deconvolution of Teh 2023 bulk RNA-seq ^10^ (n=295) using Wang 2020 scRNA-seq signature matrix: per-stage mean cell-type proportions for 16 cell-type-by-phase components.

S7 Table. ASME V&V 40-inspired verification checks (model v1.0). Nine checks across Stage A (biological plausibility, 5/5 passed) and Stage B (computational verification, 4/4 passed).

S8 Table. Window-of-implantation scenario battery: directional validation of hormone perturbations across six predefined scenarios (6/6 directional checks passed).

S8b Table. YAML PK/PD scenario snapshots: modified hormones and receptivity score at protocol timepoints for dienogest and IVF protocols (research prototype).

S9 Table. PK/PD directional pilot: literature-prior oral drugs versus baseline hormone trajectories (3/3 directional tests passed; simulation only, not transcriptome-validated).

S10 Table. Tier-B GEO datasets: 16-gene receptivity index AUC for datasets downloaded via GEO FTP (May 2026).

S1 File. Model source code, optimized parameters, and validation scripts.

---

**Fig 5.**
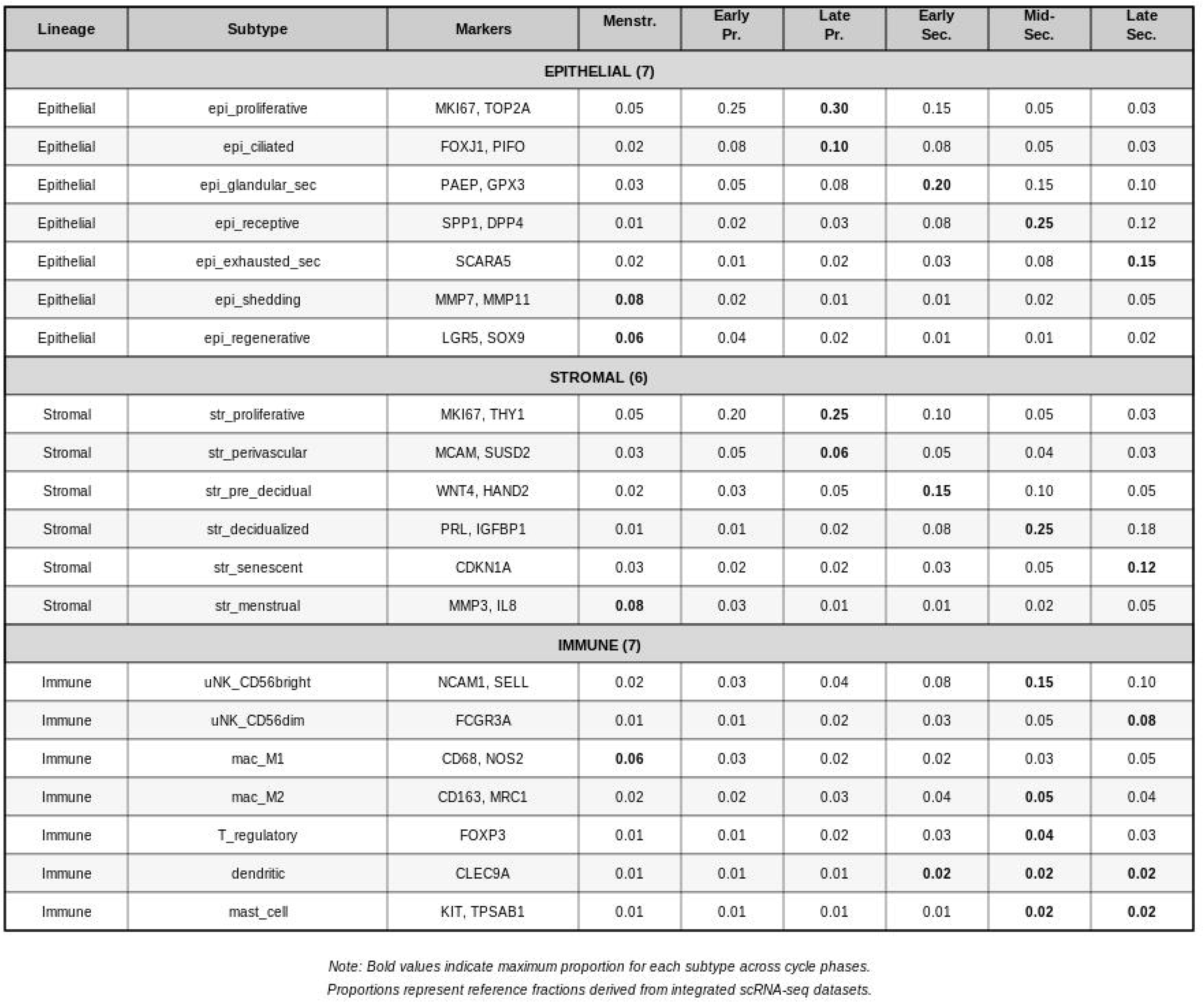
scRNA-seq cell subtypes and reference proportions. Table of 17 cell states across four compartments (epithelial, stromal, endothelial, immune) with representative marker genes and expected proportions across six menstrual cycle phases (early proliferative through late secretory), derived from Wang 2020 scRNA-seq reference data ^11^.

**Fig 6.**
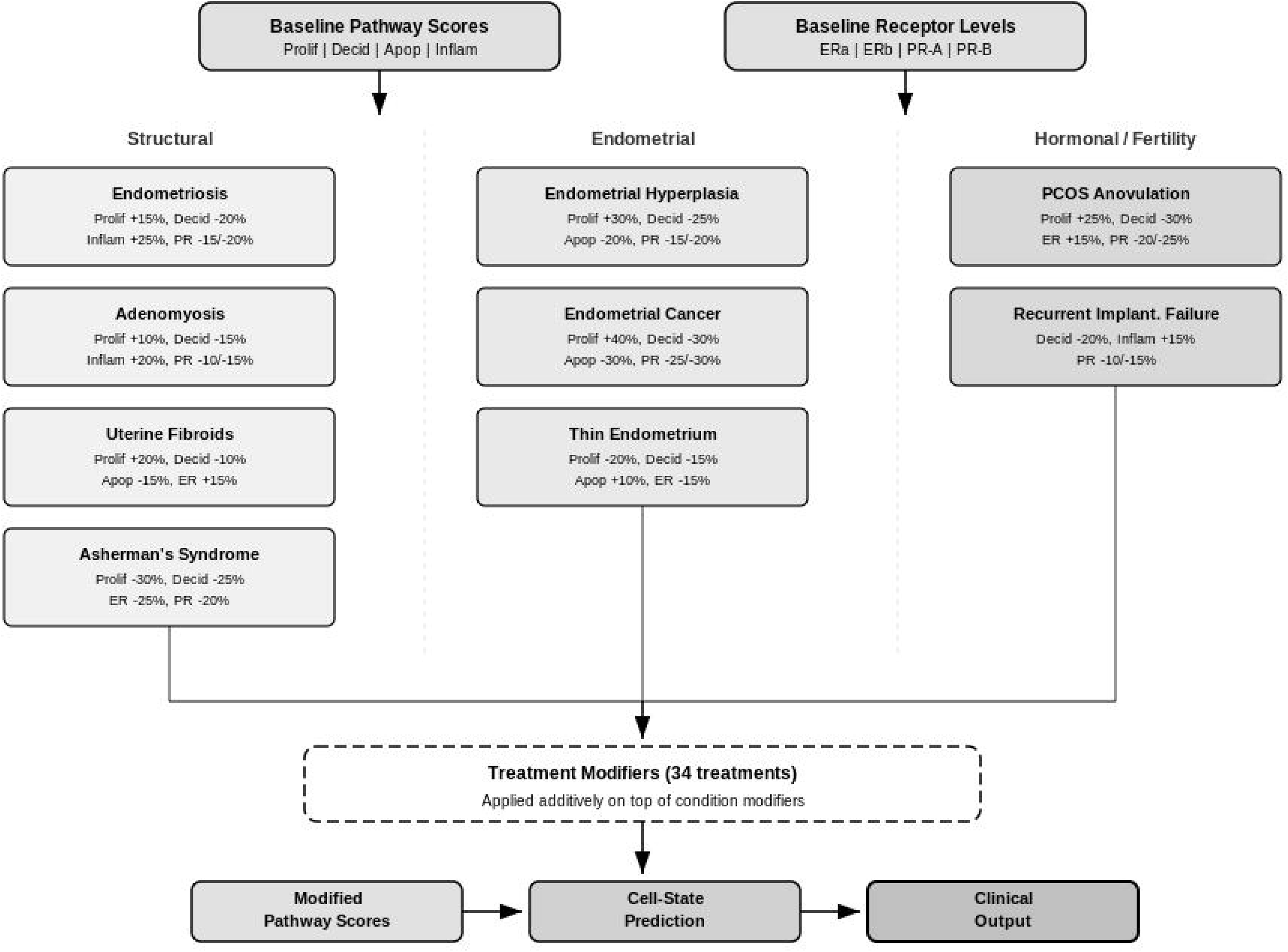
Pathological condition modifier system. Schematic showing how condition-specific disruption vectors (endometriosis, adenomyosis, PCOS, recurrent implantation failure, endometrial hyperplasia) modify baseline pathway scores and receptor levels. Three categories of modifiers are shown: structural, endometrial, and hormonal/fertility, each with condition-specific parameter adjustments validated against cross-condition transcriptomic datasets.

**Fig 7.**
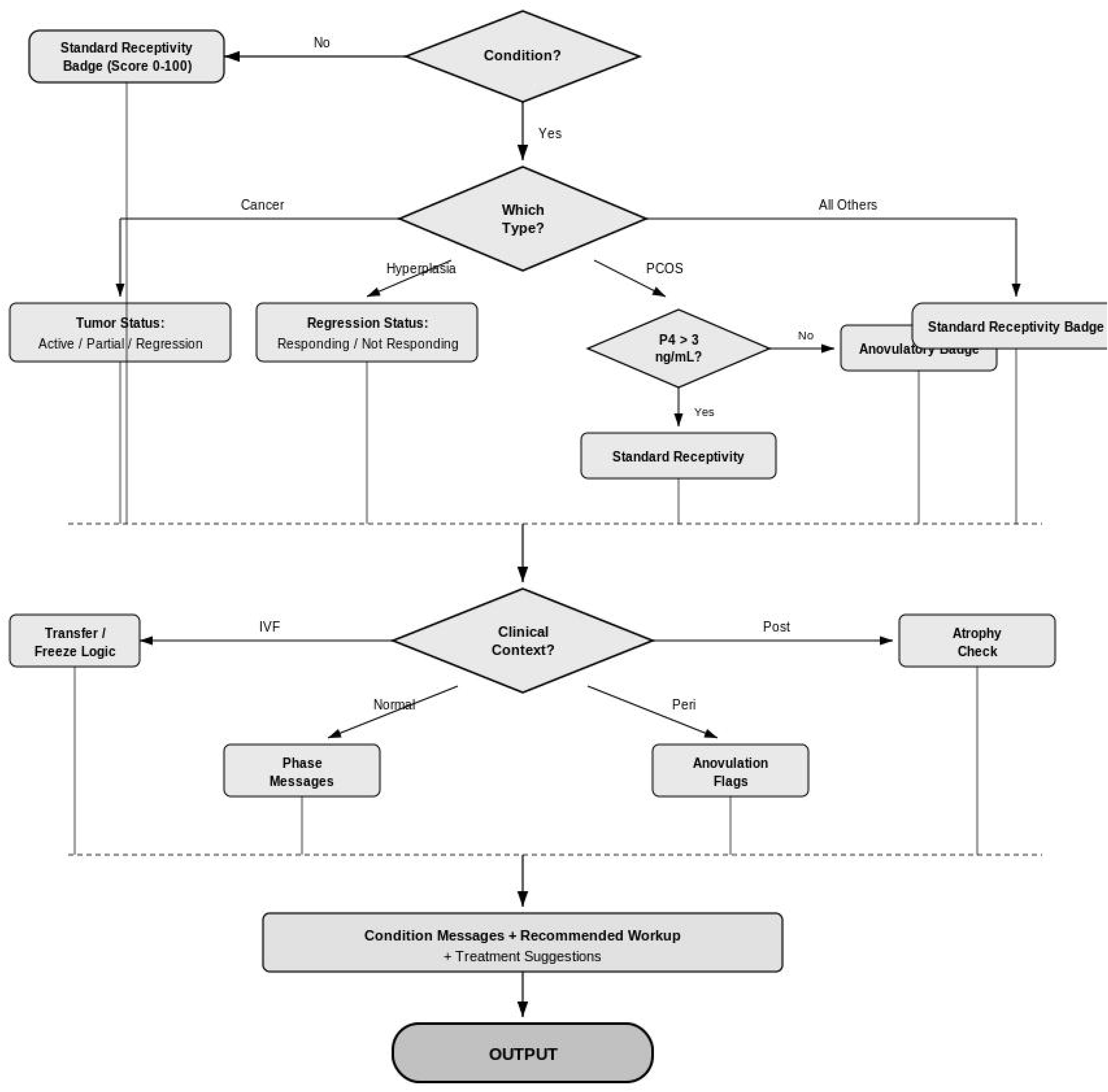
Clinical interpretation engine. Decision tree flowchart showing condition-specific clinical decision algorithms. The system accepts condition selection and biomarker inputs, computes pathway scores and Progesterone Resistance Score (PRS), and generates stratified clinical recommendations including receptivity assessment, treatment suggestions, and window of implantation timing.

**Fig 8.**
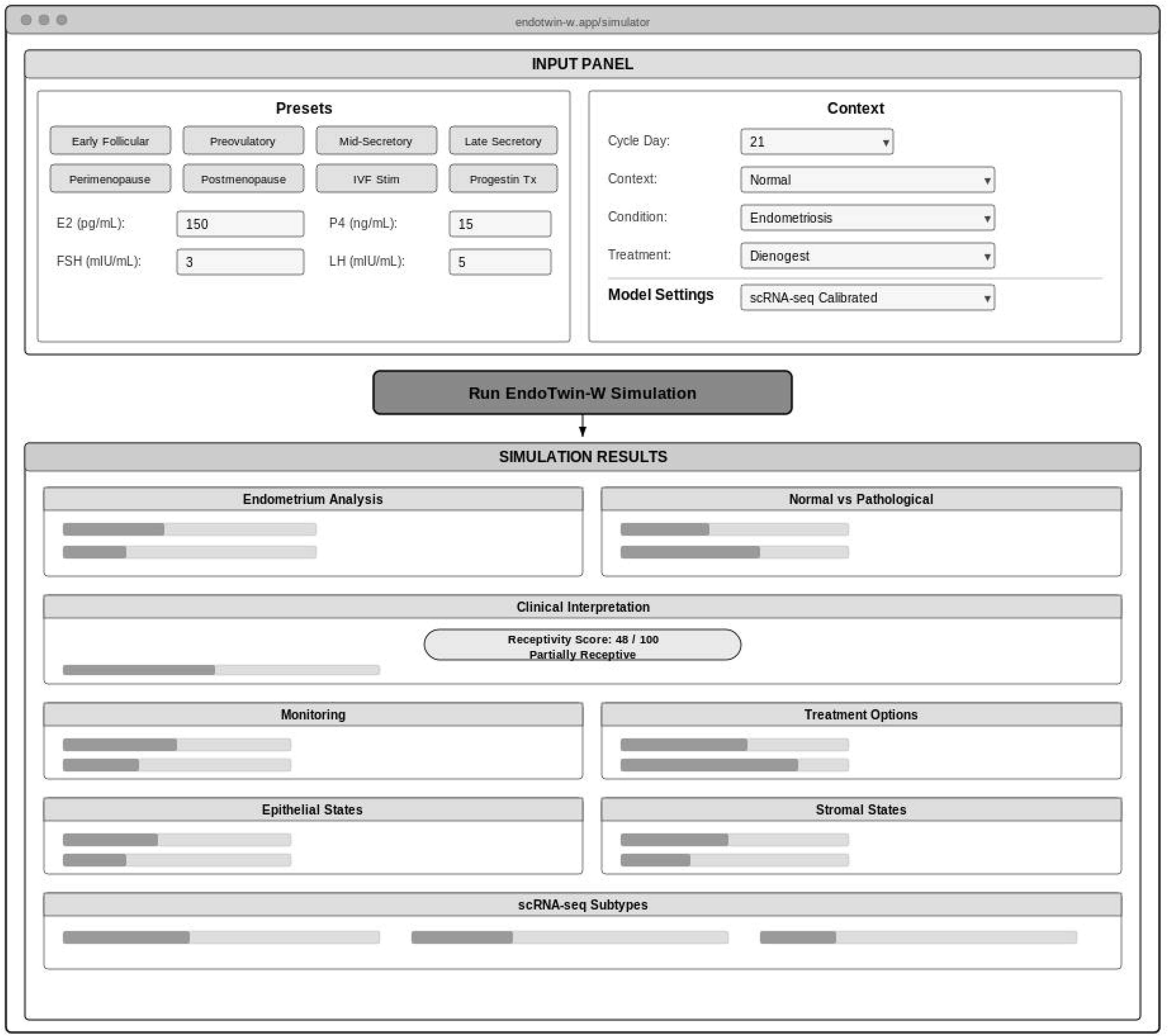
EndoTwin-W simulator interface. Screenshot of the web-based research simulator (https://endotwin-w.com) showing the user interface for condition selection, biomarker input fields, pathway score visualization, receptivity assessment output, and PDF report generation. The tool runs entirely client-side with no patient data transmitted to external servers. Outputs are for research and educational use only, not for clinical decision-making without prospective validation.

