## Supplementary material for "EndoTwin-W: glycodelin-A and CA-125 as non-invasive biomarkers of endometrial receptivity derived from a multiscale computational digital twin": File S1

### S1 File. EndoTwin-W Model Source Code, Optimized Parameters, and Validation Scripts

**Repository:**

Available from corresponding author upon request

**Web tool:**

<https://endotwin-w.com> (mirror: <https://endotwinw.com>)

**Version:** v4.0 (commit e9b831f)

**License:** All rights reserved. Code available upon request.

#### Overview

This supplementary file describes the EndoTwin-W computational model source code, optimized parameters, and validation scripts referenced in the main manuscript. All materials are available from the corresponding author upon reasonable request. The repository includes the complete Python package implementing the multiscale digital twin, validation scripts used to generate the results reported in this paper, optimized transition rate coefficients and condition-specific disruption profiles, and the marker gene reference sets used for NNLS deconvolution.

#### Repository Structure

| **Directory / File** | **Description** |
| --- | --- |
| endotwin/ | Core Python package (16 modules) |
| endotwin/model.py | Main simulation engine: ODE hormone dynamics, receptor binding, pathway scoring, CTMC state transitions |
| endotwin/hormone_dynamics.py | ODE-based estradiol and progesterone cycling with Stricker 2006 reference ranges |
| endotwin/receptor_signaling.py | Langmuir-type saturation receptor binding kinetics for ER-alpha, ER-beta, PR-A, PR-B |
| endotwin/cell_state.py | Continuous-time Markov chain (CTMC) with 17 cell states across 4 compartments |
| endotwin/condition_profiles.py | Condition-specific pathway disruption vectors (endometriosis, adenomyosis, PCOS, RIF, hyperplasia) |
| endotwin/single_cell.py | scRNA-seq calibration and NNLS deconvolution module |
| endotwin/spatial.py | Spatial compartment model: functionalis, basalis, perivascular microdomains |
| endotwin/therapeutic.py | PK/PD intervention modeling with multiplicative hormone windows |
| endotwin/woi_predictor.py | Window of implantation prediction and receptivity scoring |
| endotwin/circulating_biomarkers.py | Glycodelin-A Hill function (GdA_max=40 ng/mL, K_half=0.491, n=2.859) and CA-125 mapping |
| endotwin/phase_inference.py | Cycle phase inference from hormone levels |
| endotwin/visualize.py | Figure generation and pathway visualization |
| scripts/ | Validation and analysis scripts (27 scripts) |
| scripts/run_validation_suite.py | Master validation runner for all benchmarks |
| scripts/validate_teh2023_endotwin.py | Teh 2023 (n=236, GSE234354) external validation |
| scripts/scrna_bulk_deconvolution.py | NNLS deconvolution: 14 sub-states from 17 model states |
| scripts/cross_cohort_holdout.py | Cross-cohort held-out validation |
| scripts/cross_condition_held_out.py | Cross-condition disease validation |
| scripts/discover_markers_strict.py | Marker gene discovery (strict_markers_v34.json generation) |
| scripts/calibrate_uncertainty.py | Bootstrap uncertainty quantification |
| scripts/build_null_models.py | Null model construction for statistical comparison |
| scripts/swan_icpsr_parser.py | SWAN/ICPSR hormone data parser |
| data/ | Validation datasets and reference data |
| validation/ | Validation output logs and results |
| run_simulation.py | Command-line simulation entry point |
| requirements.txt | Python dependency list |

#### Key Parameters

The optimized transition rate coefficients (v4.0) were calibrated in a two-stage process: (1) initial fit against Wang 2020 scRNA-seq reference proportions and GSE51981 microarray data, followed by (2) refinement against NNLS-deconvolved Teh 2023 proportions using sequential Nelder-Mead/Powell optimization. The final parameter set, condition-specific disruption profiles, de-duplicated marker gene sets (strict_markers_v34.json), NNLS signature matrix, and bootstrap validation results are included in the repository.

#### Dependencies

| **Package** | **Version** |
| --- | --- |
| Python | >= 3.10 |
| NumPy | >= 1.24 |
| SciPy | >= 1.10 |
| pandas | >= 2.0 |
| scikit-learn | >= 1.2 |
| scanpy | >= 1.9 |
| anndata | >= 0.9 |
| matplotlib | >= 3.7 |
| openpyxl | >= 3.1 |

#### Reproducing Validation Results

To reproduce the validation results reported in the manuscript:

1. Clone the repository and install dependencies: pip install -r requirements.txt

2. Download the Teh 2023 dataset from GEO (GSE234354) and place in data/

3. Run the master validation suite: python scripts/run_validation_suite.py

4. Results are written to validation/ and data/ directories

The web-based research simulator at https://endotwin-w.com implements the biomarker-based pathway inference and receptivity scoring described in the manuscript. It runs entirely client-side with no data transmitted to external servers. This tool is for research use only.
